# Parvalbumin and somatostatin: biomarkers for two parallel tectothalamic pathways in the auditory midbrain

**DOI:** 10.1101/2023.08.09.552565

**Authors:** Mengting Liu, Yixiao Gao, Fengyuan Xin, Ying Hu, Tao Wang, Fenghua Xie, Tianyu Li, Ningyu Wang, Kexin Yuan

**Affiliations:** Department of Otolaryngology Head and Neck Surgery, First Affiliated Hospital of Harbin Medical University, Harbin, Heilongjiang 150001, China; Department of Basic Medical Sciences, School of Medicine, Tsinghua University, Beijing 100084, China; Tsinghua-Peking Joint Center for Life Sciences; Department of Biomedical Engineering, School of Medicine; Zhili College; School of Life Sciences; Tsinghua Laboratory of Brain and Intelligence Tsinghua University, Beijing 100084, China; Department of Otorhinolaryngology Head and Neck Surgery, Beijing Chaoyang Hospital, Capital Medical University, Beijing 100020, China; IDG/McGovern Institute for Brain Research at Tsinghua, Tsinghua University, Beijing 10084, China

## Abstract

The inferior colliculus (IC) represents a crucial relay station in the auditory pathway, located in the midbrain’s tectum and primarily projecting to the thalamus. Despite the identification of distinct cell types based on various biomarkers in the IC, their specific contributions to the organization of auditory tectothalamic pathways have remained poorly understood. In this study, we demonstrate that IC neurons expressing parvalbumin (IC^PV+^) or somatostatin (IC^SOM+^) represent major, non-overlapping cell types throughout the three IC subdivisions. Strikingly, regardless of their location within the IC, these neurons predominantly project to the primary and secondary auditory thalamic nuclei, respectively. Retrograde tracing data specific to cell types indicate that IC^PV+^ neurons primarily receive auditory inputs, whereas IC^SOM+^ neurons integrate polymodal inputs that hold behavioral significance. Furthermore, IC^PV+^ neurons exhibit significant heterogeneity in both intrinsic electrophysiological properties and presynaptic terminal size compared to IC^SOM+^ neurons. Notably, approximately one quarter of IC^PV+^ neurons are inhibitory neurons, whereas all IC^SOM+^ neurons are excitatory neurons. Collectively, our findings suggest that parvalbumin and somatostatin expression in the IC can serve as biomarkers for two functionally distinct, parallel tectothalamic pathways. This discovery challenges the conventional IC subdivision-based definition of tectothalamic pathways and calls for a reassessment of their functional roles.

## Introduction

In mammalian brains, ascending sensory pathways play critical roles in coding and relaying sensory information, changing internal states, and modulating behaviors (*Park and Friston, 2013; Choi et al., 2020; Pardi et al., 2020*). In the auditory system, nearly all auditory information ascending to the forebrain passes through the auditory midbrain, also known as the inferior colliculus (IC), making it a major station in the auditory pathway (*Batra and Fitzpatrick, 2002; Malmierca et al., 2005; Saldaña et al., 2009*). The IC consists of three subdivisions and diverse cell types and plays a critical role in various auditory brain functions, such as vocal communication (*Lohse et al., 2020*), sound localization (*Grothe et al., 2010*), and sound-induced innate behaviors (*Xiong et al., 2015; Wang et al., 2023*), that are important for animal’s survival and thrival. Given that the thalamus is the major higher brain region targeted by the IC outputs, determining how the auditory tectothalamic pathways are organized would definitely facilitate the interrogation of neural circuit mechanisms underlying the above auditory brain functions.

Our current understanding of the auditory tectothalamic pathways has been mainly based on three traditionally defined IC subdivisions, including the central nucleus (ICC), dorsal cortex (ICD) and external cortex (ICE), which were defined by cytochrome oxidase (CO) histochemistry staining (*Cant and Benson, 2005*), and based on the tectothalamic connectivity patterns revealed by traditional retrograde tracers (*Calford and Aitkin, 1983; Mellott et al., 2014; Clarke and Lee, 2018*). The ICC provides driver inputs to the ventral division of the medial geniculate body (MGBv) (*Lee and Sherman, 2010*). This pathway is regarded as the primary auditory tectothalamic pathway and required for relaying sound feature-related information to the forebrain (*Grothe et al., 2010*). The ICD and ICE provide modulatory inputs to the dorsal and medial division of the MGB (MGBd and MGBm), respectively, serving as the secondary auditory tectothalamic pathways (*Lee and Sherman, 2010*), which play important roles in mediating salient sound-induced arousal (*Wang et al., 2023*) and encoding sound detection behaviors (*Tai-Ying et al., 2023*). However, the above three auditory tectothalamic pathways defined by traditional retrograde tracers may be oversimplified. When anterograde tracers are injected into each individual subdivisions of the IC, axonal fibers from the IC can be observed in all subdivisions of the MGB, and the density of fibers in each thalamic subdivision is not negligible (*Ledoux et al., 1987; Cai et al., 2019*). In addition, abundant axonal fibers can also be observed in thalamic nuclei surrounding the MGB, such as the posterior intralaminar nucleus (PIN) (*Cai et al., 2019*) and posterior limiting nucleus (POL) (*Ledoux et al., 1987*). These data call for new approaches to define auditory tectothalamic pathways in a more realistic way.

Defining neural pathways based on the expression of specific biomarkers has been increasingly used in the neuroscience field (*Zeng, 2022*). Neurons expressing different biomarkers in the superior colliculus (SC) project to largely different downstream targets (*Shang et al., 2015; Sans-Dublanc et al., 2021; Liu et al., 2022*), for example, parvalbumin-expressing (PV+) neurons and neurotensin receptor-expressing (NTSR+) neurons project to the parabigeminal nucleus (*Shang et al., 2015*) and the paralaminar nuclei of the thalamus (*Sans-Dublanc et al., 2021*), respectively. In fact, neurons expressing different biomarkers such as parvalbumin (*Fujimoto et al., 2017*), somatostatin (*Wynne et al., 1995*), cholecystokinin (*Kreeger et al., 2021*) and vasoactive intestinal peptide (VIP) (*Goyer et al., 2019*) have also been reported in the IC, and that they are distributed throughout the IC rather than localized in an individual subdivision. Whether and how the neurons in the IC would demonstrate preference for downstream targets depending on their biomarker remained unknown.

Here, by using transgenic mice, cutting-edge viral tools, immunostaining, our newly developed rapid and deformation-free on-slide tissue-clearing method and slice recording, we identified PV+ (IC^PV+^, 14% of IC cells) and somatostatin-expressing (IC^SOM+^, 25% of IC cells) neurons as two of the major cell types in the IC that are not overlapping with each other and present in all three IC subdivisions. Interestingly, no matter in which IC subdivision the PV+ neurons are located, their axon terminals were mainly distributed in the MGBv, the primary auditory thalamus module that only projects to the primary auditory cortex. In contrast, IC^SOM+^ axon terminals are predominantly distributed in the secondary auditory thalamus, the POL in particular, which mainly projects to the posterior tail of the striatum (TS) followed by the secondary auditory cortex. We also examined the inputs of these two cell types and found that IC^PV+^ neurons primarily receive auditory inputs, whereas IC^SOM+^ neurons integrate polymodal inputs that likely hold behavioral significance. Furthermore, IC^PV+^ neurons exhibit significant heterogeneity in both intrinsic electrophysiological properties and presynaptic terminal size compared to IC^SOM+^ neurons. Notably, approximately one quarter of IC^PV+^ neurons are inhibitory neurons, whereas all IC^SOM+^ neurons are excitatory neurons. Collectively, our findings indicate that parvalbumin and somatostatin in the IC can serve as biomarkers for two parallel tectothalamic pathways, which have distinct anatomical organization pattern and functional implication. Importantly, our findings call for a revisit of functional conclusions made based on traditional definition of auditory tectothalamic pathways.

## Results

### PV+ and SOM+ neurons are two of the major cell types present in all IC subdivisions

PV+ and SOM+ neurons have been widely observed and extensively studied in many cortical and subcortical regions, and they usually play distinct roles in sensory processing and animal behaviors (*Shang et al., 2015; Tremblay et al., 2016; Anderson et al., 2018; Antonoudiou et al., 2020; Shen et al., 2022*). As mentioned earlier, these two types of neurons are present in the IC as well, but their connectivity pattern remained unknown. Therefore, we decided to focus on these two cell types in present study.

We first injected AAV-sparse-CSSP-RFP-8E3 into the IC of SOM-IRES-Cre or PV-IRES-Cre mice to characterize the morphology of sparsely labeled PV+ and SOM+ neurons. We found that both types of neurons were stellate neurons, which typically extended their dendrites beyond the single fibro-dendritic lamina (*Oliver, 2005*) (*supplement figure 1A*; SOM+: n=34 neurons from N=3 mice; PV+: n=25 neurons from N=3 mice). We next determined the distribution of PV+ and SOM+ neurons in the IC. To this end, we performed anti-NeuN immunostaining using brain slices from SOM- IRES-Cre×Ai14 mice to observe IC^SOM+^ neurons (*Figure 1A, 1^st^ column*). Although SOM-IRES-Cre mouse line has been widely used (*Taniguchi et al., 2011; Muñoz-Castañeda et al., 2021*), we were unable to validate IC^SOM+^ neurons using anti-SOM immunostaining approach because antibodies tested did not work for unknown reason. Regarding PV+ neurons, we took anti-PV immunostaining approach instead using wild-type (WT) mice (*Figure 1A, 2^nd^ column*) because very dense PV+ neurites are present in the ICC (*supplement figure 1B*), hindering the observation of PV+ cell body. We performed NeuN immunostaining using both SOM-IRES-Cre×Ai14 and WT mice to count neurons in the IC.

**Figure 1.**
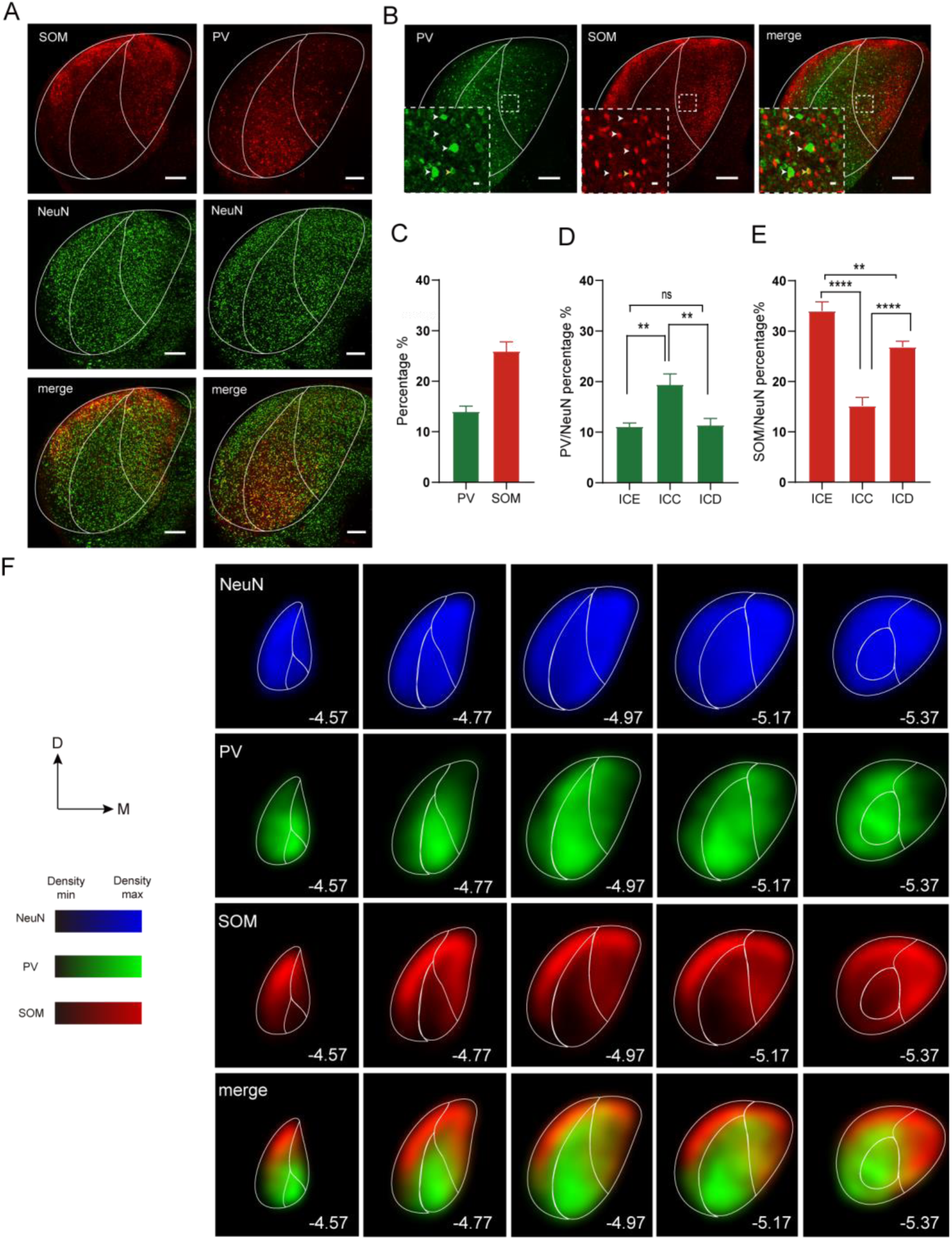
PV+ and SOM+ neurons are two of the major cell types present in all IC subdivisions. (A) Confocal 20× showing distribution of SOM+ neurons (1^st^ column, top, red), NeuN staining (1^st^ column, middle, green), an overlay (1^st^ column, bottom) in the IC, and PV+ neurons (2^nd^ column, top, red), NeuN staining (2^nd^ column, middle, green), and an overlay (2^nd^ column, bottom) in the IC. Scale bars, 200μm. (B) Confocal 20× showing PV+ neurons (left, green), SOM+ neurons (middle, red) and an overlay (right) in the IC. Scale bar, 200μm. Insets, enlarge view of the dotted box, and white arrows mark IC neurons expressing SOM or PV, while the yellow arrows mark IC neurons expressing both SOM and PV. Neurons expressing PV and SOM were mutually exclusive in the IC (right). Scale bar, 10μm. (C) Percentage of PV+ neurons (left bar, green) and SOM+ neurons (right bar, red) in the IC. n=8 slices from N=4 mice. (D) The percentages of PV+ neurons in different subdivisions of the IC (ICE, ICC and ICD). n=8 slices from N=4 mice; One-way ANOVA with post-hoc, **p<0.01, ^ns^p>0.5. (E) The percentages of SOM+ neurons in different subdivisions of the IC (ICE, ICC and ICD). n=8 slices from N=4 mice; One-way ANOVA with post-hoc, **p<0.01, ****p<0.0001; Data are means± S.E.M. (F) Kernel density estimation of neurons in the IC expressing NeuN (1^st^ row, blue), PV (2^nd^ row, green), SOM (3^rd^ row, red) and an overlay of SOM and PV (4^th^ row) from rostral (AP=-4.57) to caudal (AP=-5.37) coronal slices with a spacing of 200μm; n=20 slices from N=4 mice per group; PV (parvalbumin), SOM (somatostatin), ICC, ICD, ICE (central nucleus, dorsal cortex and external cortex of the inferior colliculus).

By performing anti-PV immunostaining using SOM-Ai14 slices, we found that neurons expressing these two biomarkers were mutually exclusive (*Figure 1B and supplement figure 1F*; n=6 slices from N=2 mice; fraction of co-expression, 2.3±0.2% of SOM+ neurons, 4.4±0.3% of PV+ neurons). Therefore, PV+ and SOM+ neurons are two distinct types of neurons in the IC in terms of the biomarker expressed.

We found that both PV+ and SOM+ neurons were distributed across all three subdivisions of the IC and in all directions (*supplement figure 1C*), and they represented 14.0±1.0% and 26.0±1.9% of neurons counted, respectively (*Figure 1C*; n=8 slices from N=4 mice), indicating that they are two of the major cell types in the IC. In terms of specific subdivisions, PV+ neurons represent about 19%, 11% and 12% of counted neurons in the ICC, ICE and ICD, respectively (*Figure 1D*; n=8 slices from N=4 mice; One-way ANOVA with post-hoc, **p<0.01), while SOM+ neurons were significantly more populated in the ICE (∼34%) and ICD (∼27%) than in the ICC (∼15%) (*Figure 1E*; n=8 slices from N=4 mice; One-way ANOVA with post-hoc, **p<0.01, ****p<0.0001). Furthermore, the distribution of these two types of neurons did not vary along the rostro-caudal axis of the IC (*supplement figures 1D and E*; n=8 slices from N=4 mice per group; unpaired t-test, p>0.5).

To more precisely determine the distribution pattern of PV+ or SOM+ neurons in the IC, we performed kernel density estimation (see Methods for more details) using five coronal slices with a spacing of 200μm (n=20 slices from N=4 mice per group). Although NeuN was evenly distributed throughout the IC (*Figure 1F, 1^st^ row*), the density of PV+ neurons decreased gradually from the ICC to ICD or ICE, while that of SOM+ neurons demonstrated an opposite decreasing direction (*Figure 1F; PV+, 2^nd^ row; SOM+, 3^rd^ row*). Collectively, the density of these two types of neurons showed a largely complementary distribution pattern, but they were comparable around the borders between ICC and ICD or ICE (*Figure 1F, 4^th^ row*).

### IC^PV+^ or IC^SOM+^ axon terminals predominantly innervate the primary or secondary auditory thalamus in an IC subdivision-independent manner

We then injected AAV-Flex-synaptophysin-mRuby into the IC of PV-IRES-Cre or SOM-IRES-Cre mice to label the synaptic terminals of PV+ or SOM+ neurons in a Cre-dependent manner (*Figure 2A*). Interestingly, in PV-IRES-Cre mice, the vast majority of PV+ synaptic terminals were localized in the ipsilateral MGBv (*Figure 2B, second row*; *supplement figure 2A*), which mainly comprises of bushy cells (*supplement figure 2C, left panel*), even when the virus was injected in a subdivision-independent manner (*Figure 2B, first row*), and only very few terminals were observed in the contralateral MGBv, the ipsilateral periaqueductal gray (PAG), the medial part of the superior olivary complex (SOCm) and the SC (*supplement figure 2D*). In contrast, SOM+ synaptic terminals were mainly distributed in the ipsilateral POL (*Figure 2C, second row*; *supplement figure 2B*), which mainly comprises of bipolar cells (*supplement figure 2C, right panel*), when virus transduction was observed in all three subdivisions (*Figure 2C, first row*), and were very sparsely located in the lateral part of ipsilateral SOC (SOCl) and in the SC (*supplement figure 2E*). No mRuby-labeled synaptic terminals was observed in the thalamus when the same virus, mixed with CTB-647 to demonstrate injection sites, was injected into the IC of WT mice (*supplement figure 2F, first row*), validating the essential dependency of mRuby expression on Cre (*supplement figure 2F, second row;* n=4 sides from N=2 mice).

**Figure 2.**
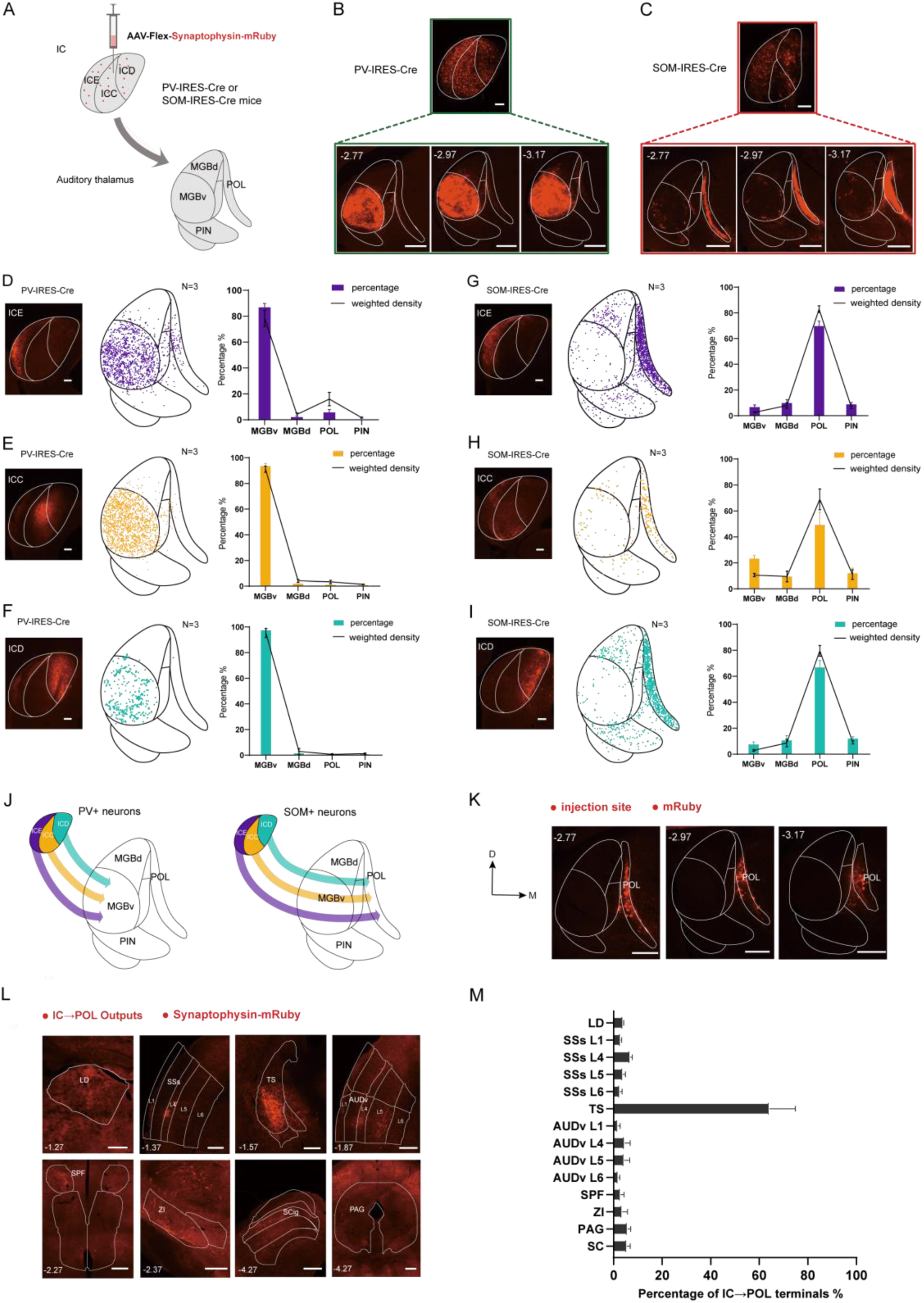
IC^PV+^ or IC^SOM+^ axon terminals predominantly innervate the primary or secondary auditory thalamus in an IC subdivision-independent manner. (A) Experimental procedures for cell type-specific anterograde tracing. Upper panel, injection of AAV-Flex-synaptophysin-mRuby into the IC of PV-IRES-Cre or SOM-IRES-Cre mice. Lower panel, anterograde projections from the IC to the major subdivisions of the auditory thalamus. (B) Example images showing when the virus was injected in a subdivision-independent manner into the IC of PV-IRES-Cre (upper green box) or SOM-IRES-Cre (upper red box), scale bars=200μm, the projections to the auditory thalamus (PV+: lower green box; SOM+: lower red box) in three AP distance (AP=-2.77, -2.97, -3.17), Scale bar=300μm. (D-F) Left panel: Example images showing when the virus was injected into the ICE (D), ICC (E) or ICD (F) of the PV-IRES-Cre mice, Scale bars=200μm. Middle panel: Distribution of IC^PV+^ terminals to the auditory thalamus sampling of 3000 terminals from each mouse brain slice (N=3) at the -2.97mm coordinate. Right panel: Percentage (bar chart) and weighted density (line chart) of IC^PV+^ neurons projecting to different subdivisions in the auditory thalamus, N=3 per group. (G-I) Left panel: Example images showing when the virus was injected into the ICE (G), ICC (H) and ICD (I) of the SOM-IRES-Cre mice, Scale bars=200μm. Middle panel: Distribution of IC^SOM+^ terminals to the auditory thalamus sampling of 300 (H) or 3000 (G, I) terminals from each mouse brain slice (N=3) at the -2.97mm coordinate. Right panel: Percentage (bar chart) and weighted density (line chart) of IC^SOM+^ neurons projecting to different subdivisions in the auditory thalamus, N=3 per group. (J) Schematic diagram illustration the primary projection of PV+ or SOM+ neurons in individual subdivisions in the IC to the auditory thalamus. PV+ neurons (left panel) in the ICE (purple), ICC (ochre) or ICD (cyan) predominately project to the MGBv. SOM+ neurons (right panel) in the ICE, ICC or ICD predominately project to the POL. (K) Example images showing expression of mRuby localized in the POL neurons, Scale bars=300μm. (L) Example images showing several target brain regions of the synaptic terminals of POL neurons receiving IC inputs, Scale bars=300μm. (M) Normalized outputs of the synaptic terminals of POL neurons receiving IC inputs in several brain regions. N=4 mice; Data are means± S.E.M.

To characterize the distribution of PV+ or SOM+ synaptic terminals in the thalamus originated from individual IC subdivisions, we injected AAV-Flex-synaptophysin-mRuby into each individual IC subdivisions of PV-IRES-Cre or SOM-IRES-Cre mice as localized as possible (N=3 mice per group). We found that, regardless of the subdivision injected, the majority of PV+ terminals were observed in the MGBv (*Figure 2E*, ICC: 94%; *Figure 2F*, ICD: 97%; *Figure 2D*, ICE: 87%; N=3 per group), while SOM+ terminals following localized virus injection in IC subdivisions were predominantly located in the secondary auditory thalamic nuclei surrounding the MGBv, in which the POL had the most axon terminals (*Figure 2G*, ICE: 70%; *Figure 2I*, ICD: 67%; *Figure 2H*, ICC: 50%; N=3 per group), followed by the MGBd and PIN. These results were consistent with the phenomenon observed following subdivision-independent virus injections (*Figures 2B and 2C*) and demonstrated a strong preference of IC^PV+^ and IC^SOM+^ projections for the primary and secondary auditory thalamus, respectively (*Figure 2J*).

Since the connectivity of the POL remained poorly understood as a subdivision of the auditory thalamus (*Márquez-Legorreta et al., 2016*), we intended to determine the downstream target regions of POL neurons receiving IC^SOM+^ inputs. Given that we had no access to the viral tool that can enable cell type-specific transsynaptic anterograde labeling, we estimated the downstream regions by injecting scAAV2/1-Cre-PA virus into the IC of WT mice to trans-synaptically express Cre in thalamic neurons receiving IC inputs, followed by targeted injections of AAV2/9-Flex-Synaptophysin-mRuby into the POL to enable Cre-dependent expression of mRuby in POL terminals (*supplement figure 2G*). Only the cases exhibiting local virus transduction in the POL (*Figure 2K*, injection site; N=4 mice) were used for further analysis. In addition, we revealed the downstream target region of the MGBv by using similar tracing strategy except that AAV2/5-Flex-tdTomoto-T2A-Synaptophysin-EGFP was locally injected into the MGBv (*supplement figures 2H and 2I*, injection site; N=4 mice).

We found that about 70% of the synaptic terminals of POL neurons receiving IC inputs (^IC^POL) were concentrated in the ipsilateral TS, followed by the AUDv (∼12%) and secondary supplemental somatosensory area (SSs) (∼12%) (*Figures 2L and 2M*). Only sparse terminals were observed in the lateral dorsal nucleus of thalamus (LD), subparafascicular nucleus (SPF), zona incerta (ZI), SC, and the PAG (*Figure 2L*). In contrast, the terminals of MGBv neurons receiving IC inputs (^IC^MGBv) were exclusively distributed in the layer 4 of AUDp (*supplement figure 2J*), which was consistent with the classic IC→MGBv→A1 circuit model established by traditional tracing method (*Ehret and Romand, 1997*). These data suggested that the major downstream target of POL neurons receiving IC^SOM+^ inputs is the TS, and that the IC^SOM+^→POL circuit belongs to secondary auditory pathway because ^IC^POL neurons only project to the secondary auditory and somatosensory cortices.

### IC^SOM+^ neurons receive significantly more polymodal inputs compared to IC^PV+^ neurons

The above data showed that IC^PV+^ and IC^SOM+^ neurons project to distinct thalamic nuclei, would they receive inputs from distinct brain regions as well? To address this question, we adopted a widely used rabies virus (RV)-mediated retrograde monosynaptic tracing strategy (*Callaway and Luo, 2015*) (see Methods for more details). Since the number of input neurons and starter neurons was linearly correlated (*Figures 3B and 3C, supplement figure 3C*), we determined the input strength of a specific region by dividing the number of input neurons in that region by the total number of input neurons (N=4 mice per group).

**Figure 3.**
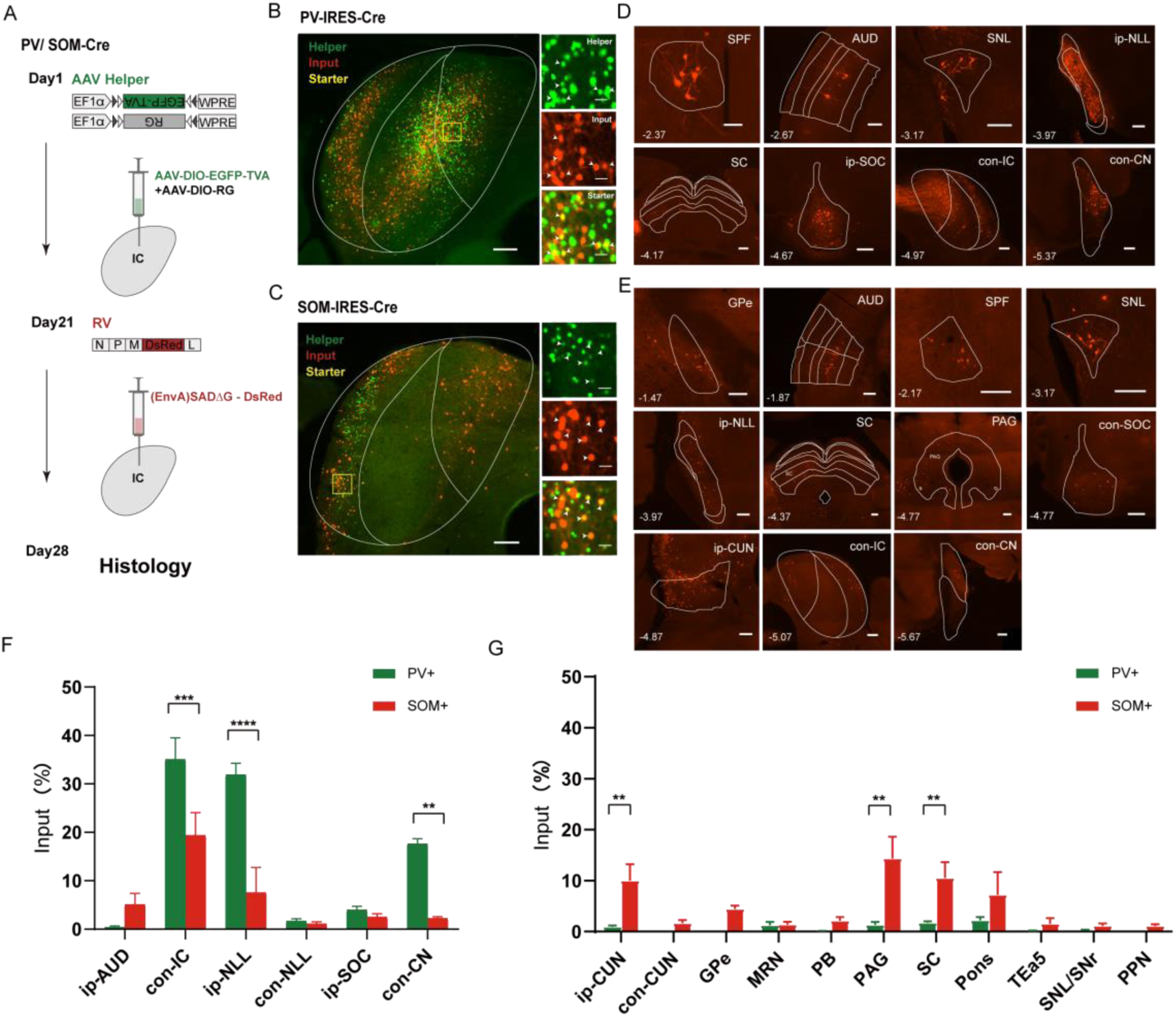
IC^SOM+^ neurons receive significantly more polymodal inputs compared to IC^PV+^ neurons. (A) Schematic diagram showing RV-mediated monosynaptic retrograde tracing. Left, time course of virus injection and histology. Right, construction of AAV helper viruses (AAV-DIO-EGFP-TVA and AAV-DIO-RG) and pseudotyped RV. (B-C) Left panel, representative fluorescent images showing the infection of RV and AAV in the IC of PV-IRES-Cre (B) or SOM-IRES-Cre (C) mice, scale bars=200μm. Right panel, enlarged view of boxed area showing the expression of AAV-Helper, input and starter, respectively, scale bars=25μm. White arrows in the right panel showing starter neurons expressing both EGFP and DsRed. (D) Example images showing major inputs of the PV+ neurons in the IC, and the arrangement of the images is from the anterior to the posterior. Scale bars, 200μm. (E) Example images showing major inputs of the SOM+ neurons in the IC, and the arrangement of the images is from the anterior to the posterior. Scale bars, 200μm. (F) Normalized inputs of IC^SOM+^ and IC^PV+^ neurons from auditory-related brain regions. N=4 mice per group, two-way ANOVA with Geisser-Greenhouse correction, **p<0.01, ***p<0.001, ****p<0.0001. (G) Normalized inputs of IC^SOM+^ and IC^PV+^ neurons from polymodal-related brain regions. N=4 mice per group, two-way ANOVA with Geisser-Greenhouse correction, **p<0.01. Data are means± S.E.M.

We found that IC^PV+^ neurons received inputs predominantly from the auditory brainstem (∼91%), mainly including contralateral IC (∼35%), ipsilateral NLL (∼32%) and contralateral CN (∼18%) (*Figures 3D and 3F; supplement figure 3D and 3E*; N=4 mice per group, two-way ANOVA with Geisser-Greenhouse correction), suggesting that IC^PV+^ neurons belong to the classic ascending auditory pathway. In contrast, IC^SOM+^ neurons received significantly more inputs from polymodal sensorimotor regions (∼55%), including the PAG (∼14%), SC (∼11%) and CUN (∼11%) (*Figures 3E and 3G; supplement figures 3D and 3E*; N=4 mice per group, two-way ANOVA with Geisser-Greenhouse correction), than from auditory-related brain regions (∼38%), mainly including the contralateral IC (∼18%) and ipsilateral NLL (∼7%) (*Figure 2F, supplement figures 3D and 3E*). Interestingly, IC^SOM+^ neurons received very few inputs from contralateral CN (*Figures 3E and 3F*), further indicating that these neurons do not belong to the primary auditory pathway.

Together with the outputs of IC^PV+^ and IC^SOM+^ neurons, these data suggest that IC^PV+^ neurons may be mainly responsible for relaying sound feature-related information from the peripheral to the auditory cortex via the primary auditory thalamus, while IC^SOM+^ neurons can integrate polymodal, behaviorally meaningful inputs to modulate mice’s behavioral states via the POL→TS pathway.

### IC^PV+^ neurons are anatomically more heterogenous than IC^SOM+^ neurons

The distinct input-output architectures of IC^SOM+^ and IC^PV+^ neurons inspired us to hypothesize that these two types of neurons may have distinct anatomical features, which could contribute to the potential functional distinctions between them.

To test the above hypothesis, we first performed anti-GAD and anti-PV immunostaining using brain slices containing the IC from SOM-IRES-Cre×Ai14 and VGAT×Ai14 mice, respectively. GAD and VGAT, which are commonly used markers for GABAergic neurons (*Ono et al., 2005; Ito et al., 2009; Oberle et al., 2023*), are co-expressed in IC neurons (*supplement figure 4A*). We found that, in all IC subdivisions, no SOM+ neurons were labeled by anti-GAD immunostaining (*Figure 4A*; n=12 slices from N=3 mice), indicating that all SOM+ neurons in the IC are excitatory. In contrast, about 25% of counted PV+ neurons were VGAT+, and this phenomenon was IC subdivision-independent (*Figure 4B and supplement figure 4B*; n= 12 slices from N=3 mice; ICE: 26±1.5%, ICC: 21±3%, ICD: 28±3.6%), indicating that IC^PV+^ neurons can be divided into inhibitory and excitatory groups. These observations were further confirmed by anti-GAD immunofluorescent staining of mRuby-labeled IC^SOM+^ or IC^PV+^ terminals in the thalamus, which showed that all IC^SOM+^→POL terminals were GAD-whereas IC^PV+^→MGBv terminals were either GAD-or GAD+ (*Figure 4C*), and the inhibitory IC^PV+^→MGBv terminals accounted for about 12% of the total IC^PV+^→MGBv terminals (*Figure 4C, bar chart*; n=6 slices from N=3 mice).

**Figure 4.**
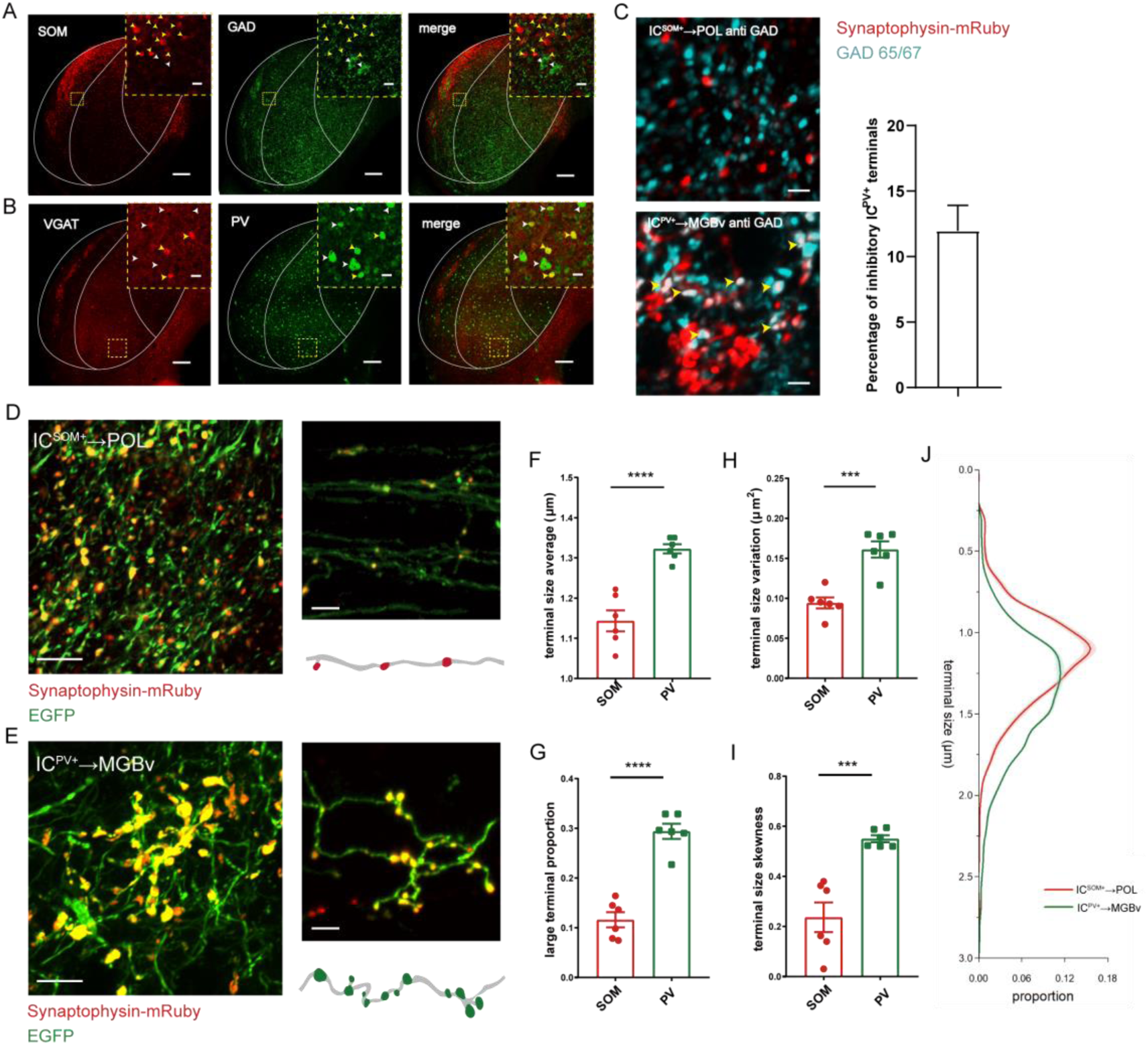
IC^PV+^ neurons are anatomically more heterogenous than IC^SOM+^ neurons. (A) Confocal 20× images showing IC^SOM+^ neurons (red, left), GAD65/67 staining (green, middle), and an overlay (right). Scale bar, 200μm. Insets, enlarge view of the dotted box, and white arrows mark IC neurons expressing GAD65/67, while the yellow arrows mark IC neurons expressing SOM. There was no overlap between SOM+ neurons and GABAergic neurons (right). Scale bar, 10μm. (B) Confocal 20× images showing IC^VGAT+^ neurons (red, left), PV staining (green, middle), and an overlay (right), Scale bars, 200μm. Insets, enlarge view of the dotted box, and white arrows mark IC neurons expressing PV, while the yellow arrows mark IC neurons expressing VGAT. There were some overlaps between PV+ neurons and GABAergic neurons (right), Scale bars, 10μm. (C) Confocal 40× images showing IC^SOM+^→POL terminals (upper panel; red channel: synaptophysin-mRuby), IC^PV+^→MGBv terminals (lower panel; red channel: synaptophysin-mRuby), and GAD65/67 staining (cyan channel). There was no overlap between IC^SOM+^→POL terminals and GAD65/67 (upper panel), while there were some overlaps between IC^PV+^→MGBv terminals and GAD65/67 (lower panel, yellow arrows). The bar chart on the right side showing the percentage of inhibitory IC^PV+^→MGBv terminals, n=6 slices from N=3 mice. (D-E) Confocal 40× images showing the morphological features of IC^SOM+^→POL terminals (D) and IC^PV+^→MGBv terminals (E). Scale bar, 100μm. The upper right panel showing sparer terminals by sparse labeling of IC^SOM+^ (D) and IC^PV+^ neurons (E), and the lower right panel showing a schematic diagram of the morphology of IC^SOM+^→POL (D) and IC^PV+^→MGBv terminals (E). Scale bars, 5μm. (F) Average terminal sizes of IC^SOM+^→POL (N=6 mice) and IC^PV+^→MGBv (N=6 mice). unpaired t-test, p<0.0001. (G) Large terminal (defined as terminals with sizes larger than 1.5 μm) proportion of IC^SOM+^→POL (N=6 mice) and IC^PV+^→MGBv (N=6 mice). unpaired t-test, p<0.0001. (H) Terminal size variation of IC^SOM+^→POL (N=6 mice) and IC^PV+^→MGBv (N=6 mice). unpaired t-test, p<0.001. (I) Terminal skewness of IC^SOM+^→POL (N=6 mice) and IC^PV+^→MGBv (N=6 mice). unpaired t-test, p<0.001. Data are means± S.E.M. (J) Terminal size distribution of IC^SOM+^→POL (N=6 mice) and IC^PV+^→MGBv (N=6 mice). The SEM is shown as light shadows. The bins are 0.1μm.

We next characterized the morphological features of IC^SOM+^ and IC^PV+^ terminals in the POL and MGBv, respectively, to estimate their potential influences on their postsynaptic thalamic neurons. Toward this end, we injected AAV2/9-flex-synaptophysin-mRuby into the IC of SOM-IRES-Cre or PV-IRES-Cre mice to label IC^SOM+^ or IC^PV+^ terminals and utilized our newly developed rapid and deformation-free on-slide tissue-clearing method to visualize the terminal morphology and terminal size (*supplement figure 4C*; see Methods for more details).We further used AAV2/9-flex-synaptophysin-mRuby-T2A-EGFP to confirm the localization of mRuby signals at axonal swellings suggesting presynaptic terminals, including boutons and varicosities (*Figures 4D and 4E*).

Compared with the sizes of IC^SOM+^→POL terminals, those of IC^PV+^→MGBv terminals were averagely larger (*Figures 4D, 4E and 4F*; N=6 mice per group; PV+ terminals, 1.3±0.01 μm; SOM+ terminals, 1.1±0.02 μm; unpaired t-test, p<0.0001), partly contributed by a higher proportion of terminals that were larger than 1.5 μm (*Figure 4G*; PV+ terminals, 0.29± 0.02; SOM+ terminals, 0.12±0.02; unpaired t-test, p<0.0001), and more heterogeneous (*Figure 4H*, PV+ terminals, 0.16±0.01; SOM+ terminals, 0.09±0.01; unpaired t-test, p<0.001), and their distribution was more deviated from the normal distribution (*Figure 4I*; PV+ terminals, 0.55±0.01; SOM+ terminals, 0.24±0.06; unpaired t-test, p<0.001). All of the above phenomena can also be found in the distribution diagram of the two groups’ terminal size (*Figure 4J*), which additionally showed that the representative sizes of IC^PV+^→MGBv terminals spanned from 0.9 to 1.8 μm, but those of IC^SOM+^→POL terminals were restricted to about 1.1 μm.

Taken together, the anatomical features of IC^PV+^ neurons are distinct from and more heterogeneous than IC^SOM+^ neurons.

### IC^PV+^ neurons are electrophysiologically more heterogenous than IC^SOM+^ neurons

We next conducted whole-cell current-clamp recordings from IC^PV+^ or IC^SOM+^ neurons by using slices from PV-IRES-Cre×Ai14 or SOM-IRES-Cre×Ai14 mice to characterize their intrinsic electrophysiological properties. We found that the firing of recorded IC^SOM+^ neurons (n=22 cells from N=4 mice) in response to current injection uniformly demonstrated adapting profile (*Figure 5A, 1*). With the increase of current amplitude, the firing rate of these neurons increased nearly linearly at first and then reached a plateau (*Figure 5B, red triangle*). In contrast, three types of firing profiles were observed in IC^PV+^ neurons (n=28 cells from N=4 mice), including rapid inactivating (RI; *Figure 5A, 3*; n=12 cells), regular spiking (RS; *Figure 5A, 4*; n=12 cells) and fast spiking (FS; *Figure 5A, 2;* n=4 cells). RI neurons only fired one or two action potentials at the onset of current injection, and their firing rate barely changed with the increase of current amplitude (*Figure 5B, green squares*). The firing rate of FS neurons was considerably higher than that of RS neurons. Although the firing rate of both types of neurons increased largely linearly with current amplitude, that of FS neurons increased much faster compared with RS neurons (*Figure 5B*). Thus, the firing mode of IC^PV+^ neurons was more diverse than that of IC^SOM+^ neurons.

**Figure 5.**
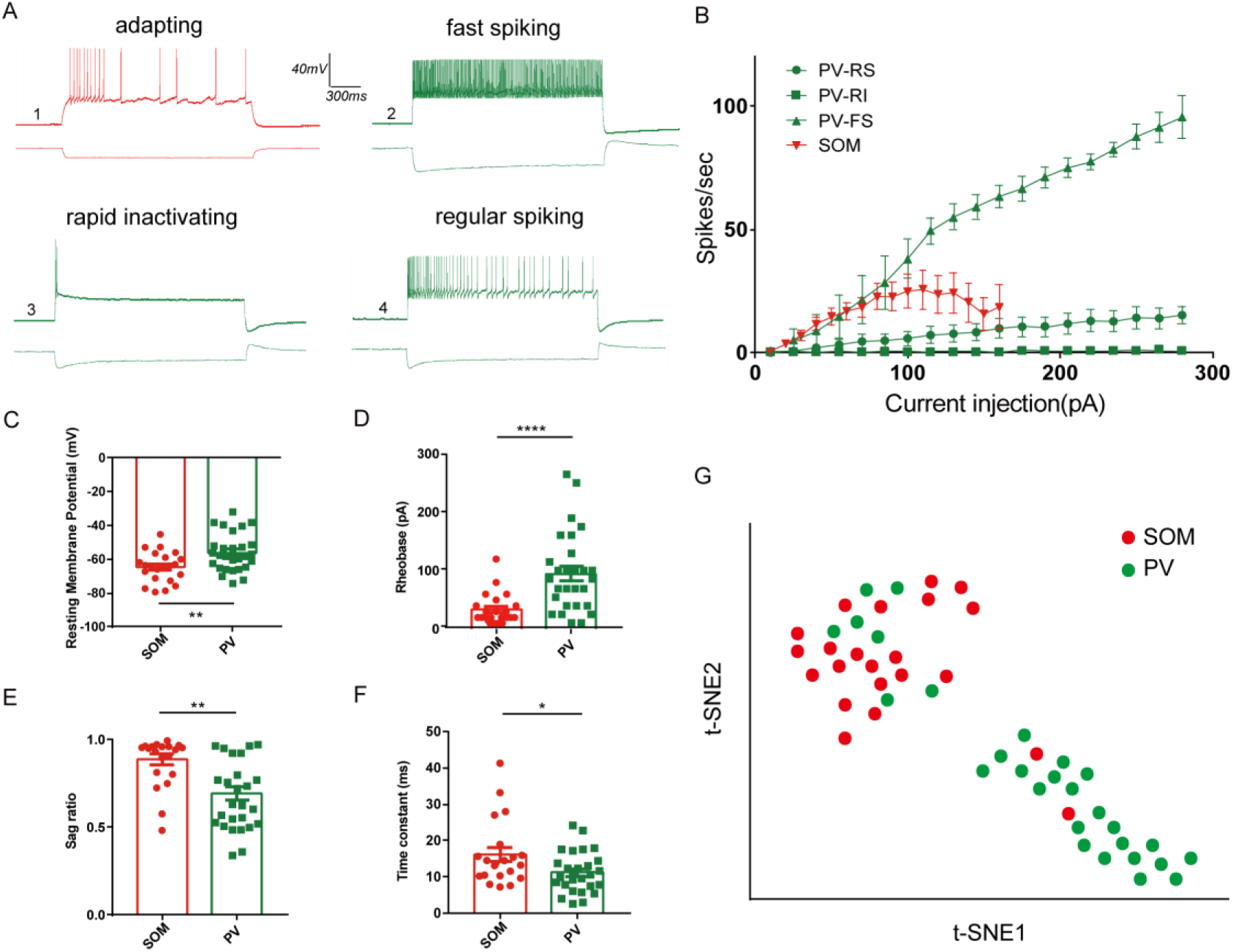
IC^PV+^ neurons are electrophysiologically more heterogenous than IC^SOM+^ neurons. (A) Representative traces from whole-cell current-clamp recordings obtained from SOM+ and PV+ neurons in the IC. (A1) shows adapting profile of IC^SOM+^ neurons, (A2) fast spiking of IC^PV+^ neurons, (A3) rapid inactivating of IC^PV+^ neurons and (A4) regular spiking of IC^PV+^ neurons. Scale bars, 40mV/300ms. (B) Mean values of step current vs. spikes/sec (I/F) curves calculated from all recorded IC^SOM+^ (red triangles, n=22 cells from N=4 mice) and IC^PV+^ neurons showing the most common spiking classes: fast (green triangles, n=4 cells from N=4 mice), regular (green circles, n=12 cells from N=4 mice) and rapid inactivating (green squares, n=12 cells from N=4 mice). (C) Resting membrane potential of IC^SOM+^ neurons (n=22 cells from N=4 mice) versus IC^PV+^ neurons (n=28 cells from N=4 mice), unpaired t test, p<0.01. (D) Rheobase of IC^SOM+^ neurons versus IC^PV+^ neurons, Mann-Whitney test, p<0.0001. (E) Sag ratio of IC^SOM+^ neurons versus IC^PV+^ neurons, Mann-Whitney test, p<0.01. (F) Time constant of IC^SOM+^ neurons versus IC^PV+^ neurons, Mann-Whitney test, p<0.05. Data are means± S.E.M. (G) Two-dimensional t-SNE representation of electrophysiological properties for two molecular cell types. Clustering was performed using electrophysiological parameters we analyzed.

We further compared the membrane properties between SOM+ and PV+ neurons. We found that, compared with IC^PV+^ neurons, IC^SOM+^ neurons had significantly lower resting membrane potential (*Figures 5C*; IC^SOM+^: -64.6±1.9 mV; IC^PV+^: -56.2±2.1 mV; unpaired t-test, p<0.01), lower rheobase (*Figures 5D*; IC^SOM+^: 33.6±5.7 pA; IC^PV+^:95.2±12.7 pA; Mann-Whitney test, p<0.0001), and higher sag ratio (*Figures 5E*; IC^SOM^: 0.9±0.03; IC^PV+^: 0.7±0.04; Mann-Whitney test, p<0.01), suggesting significant differences in the expression of potassium, sodium and hyperpolarization-activated channels, respectively, between IC^SOM+^ and IC^PV+^ neurons. We also noticed that the rheobase and sag ratio of IC^PV+^ neurons were distributed in a wide range. In addition, the time constants of IC^SOM+^ neurons were significantly higher than that of IC^PV+^ neurons (*Figure 5F*; IC^SOM^: 16.1±2.0 ms; IC^PV+^: 11.2±1.1 ms; Mann-Whitney test, p<0.05), suggesting that the soma of IC^SOM+^ neurons was larger.

Considering that IC^SOM+^ and IC^PV+^ neurons demonstrated distinctions in multiple dimensions of intrinsic electrophysiological properties, we performed clustering and dimensionality-reduction using the t-SNE algorithm to obtain a two-dimension embedding of these two cell types (*Figure 5G*). The t-SNE plot showed two well-separated clusters, one was mainly composed of IC^SOM+^ and the other one predominantly comprised IC^PV+^ neurons. The more dispersed distribution of IC^PV+^ neurons was consistent with their higher degree of heterogeneity in intrinsic electrophysiological properties.

## Discussion

The inferior colliculus (IC), which consists of three subdivisions and diverse cell types, plays a critical role in various important auditory brain functions, such as sound localization (*Masterton and Imig, 1984; Grothe et al., 2010*), auditory plasticity (*Bajo et al., 2010*), auditory contrast gain control (*Lohse et al., 2020*), integration of polymodal signals (*Chen and Song, 2019; Yang et al., 2020; Lohse et al., 2021*), sound detection (*Tai-Ying et al., 2023*), auditory arousal (*Wang et al., 2023*) and sound-induced defensive behavior (*Xiong et al., 2015; Wang et al., 2023*). Given that the thalamus is the major higher brain region targeted by the IC, investigating tectothalamic organization will greatly benefit our understanding of the way in which the IC routes and transmits information to the thalamus for further processing and interpretation, facilitating the interrogation of neural circuit mechanisms underlying the above auditory brain functions. In the present study, we showed that PV+ and SOM+ neurons in the IC, regardless of their location, predominantly contribute to the primary and secondary auditory pathways, respectively. Remarkable differences in input sources, intrinsic electrophysiological properties and axon terminal features collectively suggested that these two types of neurons are functionally distinct (*Figure 6*). Our findings reconciled the discrepancies among previous studies characterizing tectothalamic pathways based on the subdivisions of the IC and provided an anatomical framework for molecular marker-based, functional interrogation of the tectothalamic pathways.

**Figure 6.**
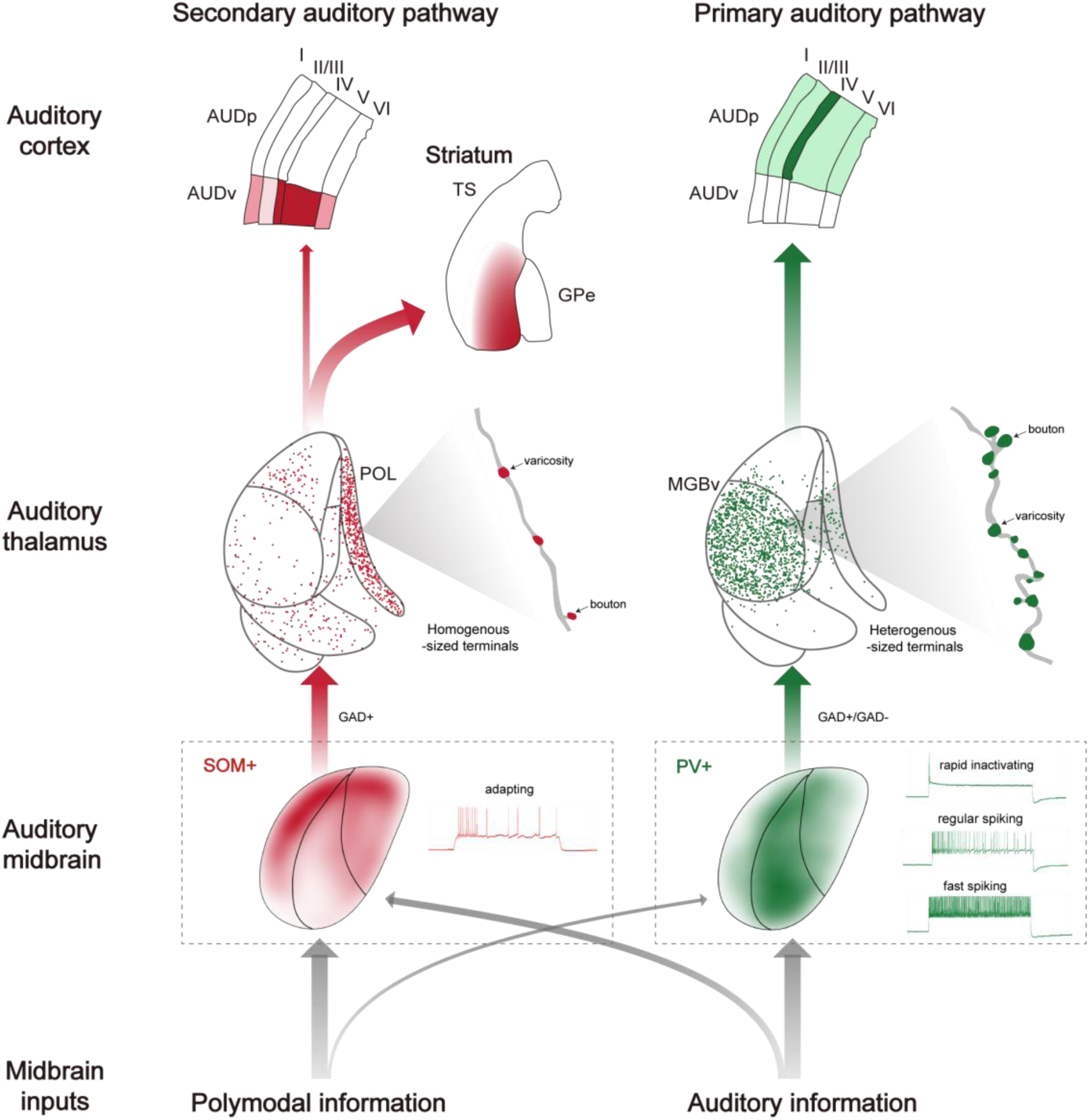
Schematic diagram illustrating the secondary and primary auditory pathways by IC^SOM+^ and IC^PV+^ neurons. The schematic diagram contains distribution, intrinsic electrophysiological property, axon terminal features and input-output architecture of IC^SOM+^ (red) and IC^PV+^(green) neurons. Darker color represents higher distribution density or stronger projections of IC^SOM+^ or IC^PV+^ neurons.

### Biomarker-based versus subdivision-based auditory tectothalamic pathways

Traditionally, auditory tectothalamic pathways have been defined based on the three subdivisions of the IC, highlighting topographic innervation of the auditory thalamus by different IC subdivisions. This definition was mainly derived from retrograde tracing studies using traditional tracers such as HRP (*Calford and Aitkin, 1983*) and retro-beads (*Mellott et al., 2014*). However, data in these studies has also shown that an individual thalamic subdivision can actually receive inputs from all IC subdivisions despite the preference for certain IC subdivision. Echoing with this observation, studies using traditional anterograde tracer show that neurons in an individual IC subdivision can send their axons to several auditory thalamic subdivision (*LeDoux et al., 1985*). These results strongly suggest that the auditory tectothalamic pathways defined by IC subdivisions may have been oversimplified. In the present study, by combining transgenic mice with cutting-edge viral tracing tools and out newly developed tissue-clearing method, we showed that PV+ and SOM+ neurons make distinct projections to the auditory thalamus in an IC subdivision-independent manner. This finding provides an alternative approach to define auditory tectothalamic pathways and can likely interpret the intricate observations made by using traditional tracers. Nevertheless, our results do not exclude the possibility that IC neurons expressing other biomarkers may also contribute to distinct tectothalamic pathways, although it has been shown that VIP+ neurons do not do so (*Beebe et al., 2022*).

### Potential function of IC^PV+^ and IC^SOM+^ neurons-mediated auditory pathways

We showed that IC^PV+^ neurons represent about 15% of IC neurons, and that receive inputs predominantly from the contralateral IC, ipsilateral NLL and contralateral CN, which are the major stations in ascending auditory pathway (*Oliver et al., 2018*), strongly suggesting that IC^PV+^ neurons mainly receive auditory inputs. We also demonstrated that IC^PV+^ neurons are highly heterogeneous in terms of their intrinsic electrophysiological properties and axon terminal size. This remarkable heterogeneity can likely enhance the robustness of the representation and transmission of time-varying acoustic features by IC^PV+^ neurons (*Chen et al., 2022*). Furthermore, the MGBv, the primary auditory thalamus, is the dominating downstream target of IC^PV+^ neurons. These data collectively suggest that IC^PV+^ neurons may play an important role in auditory processing by providing the MGBv with both feedforward excitation and inhibition.

In contrast, IC^SOM+^ neurons, which represent about 25% of IC neurons, can likely integrate auditory inputs with polymodal, threat-indicative inputs from multiple sources, such as the PAG (*Lefler et al., 2020*), SC (*Basso et al., 2021*) and CUN (*Caggiano et al., 2018*), suggesting that IC^SOM+^ activity correlates with animal behaviors. Interestingly, both the electrophysiological and anatomical heterogeneity of IC^SOM+^ neurons are much lower than those of IC^PV+^ neurons, suggesting that these neurons are more suitable for sensory detection rather than discrimination and capable of effective information transmission. Furthermore, IC^SOM+^ neurons predominantly project to the POL, which in turn mainly innervates the TS. It has been shown that TS neurons can mediate alerting stimuli-induced arousal (*Wang et al., 2023*) and the reinforcement of defensive behavior (*Menegas et al., 2018*) in freely moving mice. Together, our data suggest that IC^SOM+^ neurons are responsible for detecting polymodal, alerting stimuli and effectively triggering the transition of behavioral state to defense.

### Molecular markers as an effective approach for classifying cell types in the IC

The classification of cells into distinct types are crucial for understanding the function and dysfunction of certain brain regions. Combining gene expression patterns with neuronal connectivity has been proven to be an effective approach to define cell types (*Zeng and Sanes, 2017; Zeng, 2022*). In the present study, we showed that PV+ and SOM+ neurons in the IC are two groups of neurons that are non-overlapping and have distinct thalamic targets, likely underlying distinct functions. This observation is reminiscent of that reported in the SC (*Liu et al., 2022*). Specifically, PV+ neurons in the SC predominantly project to the parabigeminal nucleus to trigger fear responses (*Shang et al., 2015*), whereas NTSR+ neurons mainly innervate the paralaminar nuclei of the thalamus, promoting visual stimuli-induced defensive emotional state (*Sans-Dublanc et al., 2021*). Thus, in both the auditory and visual midbrains, some of the biomarkers can be used to define downstream pathways that likely play distinct functions in sensory perception.

### Technical limitations

Given that anterograde tracing virus that can enable cell type-specific transsynaptic labeling was not available to us, we were not able to selectively characterize the downstream targets of POL neurons receiving IC^SOM+^ inputs. Although the axon terminals of POL neurons are predominantly distributed in the TS, about 20% of the terminals were also observed in the AuDv and SSs. Therefore, the exact distribution pattern of the axon terminals of POL neurons receiving IC^SOM+^ input remains unclear. Nevertheless, our data showed that the axon terminals of IC^SOM+^ neurons cover the entire POL in both AP and DV directions with high density (*supplement figure 2B*), suggesting that most POL neurons receive IC^SOM+^ input. Thus, it is very likely that the projection pattern of POL neurons revealed by our cell type-independent approach applies to IC^SOM+^-targeted POL neurons as well.

## Acknowledgments

We thank B.H. for instrumental support, S. S. for SOM-IRES-Cre mice supplement, and Y. Z. for assistance of using ANDOR OXFORD Instrument (Imaging Core Facility, Technology Center for Protein Sciences, Tsinghua University). K.Y. receives funding from STI 2030-Major Projects 2021ZD0200300, National Natural Science Foundation of China (31871057, 32070993, 81527901, T2341003), Beijing Municipal Science & Technology Commission (Z181100001518004, Z181100001518006), and Guoqiang Institute, Tsinghua University. N.W. receives funding from National Natural Science Foundation of China (81770993) and Capital’s Funds for Health Improvement and Research (2020-1-2032).

## Author contributions

Conceptualization, K.Y.; Methodology and Investigation, M.L., Y.G., F.Xin, Y.H., T.W., F.Xie and T. L.; Data analysis, M.L., Y.G., F.Xin and Y.H.; Data Curation, all authors; Writing – Original Draft, K.Y., M.L. and Y.G.; Writing – Review & Editing, K.Y., M.L. and Y.G.; Supervision, K.Y. and N.W.

## Materials and methods

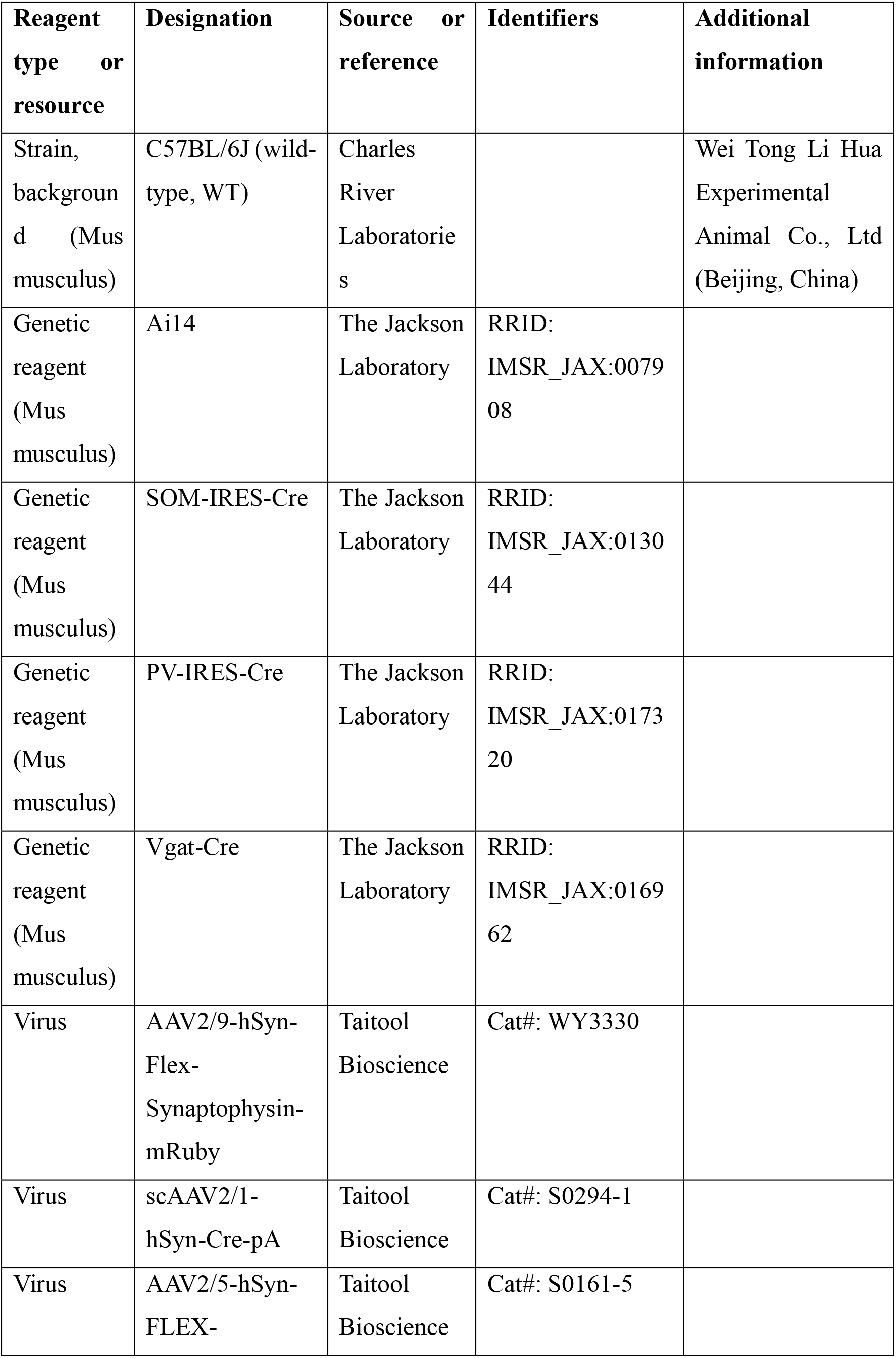
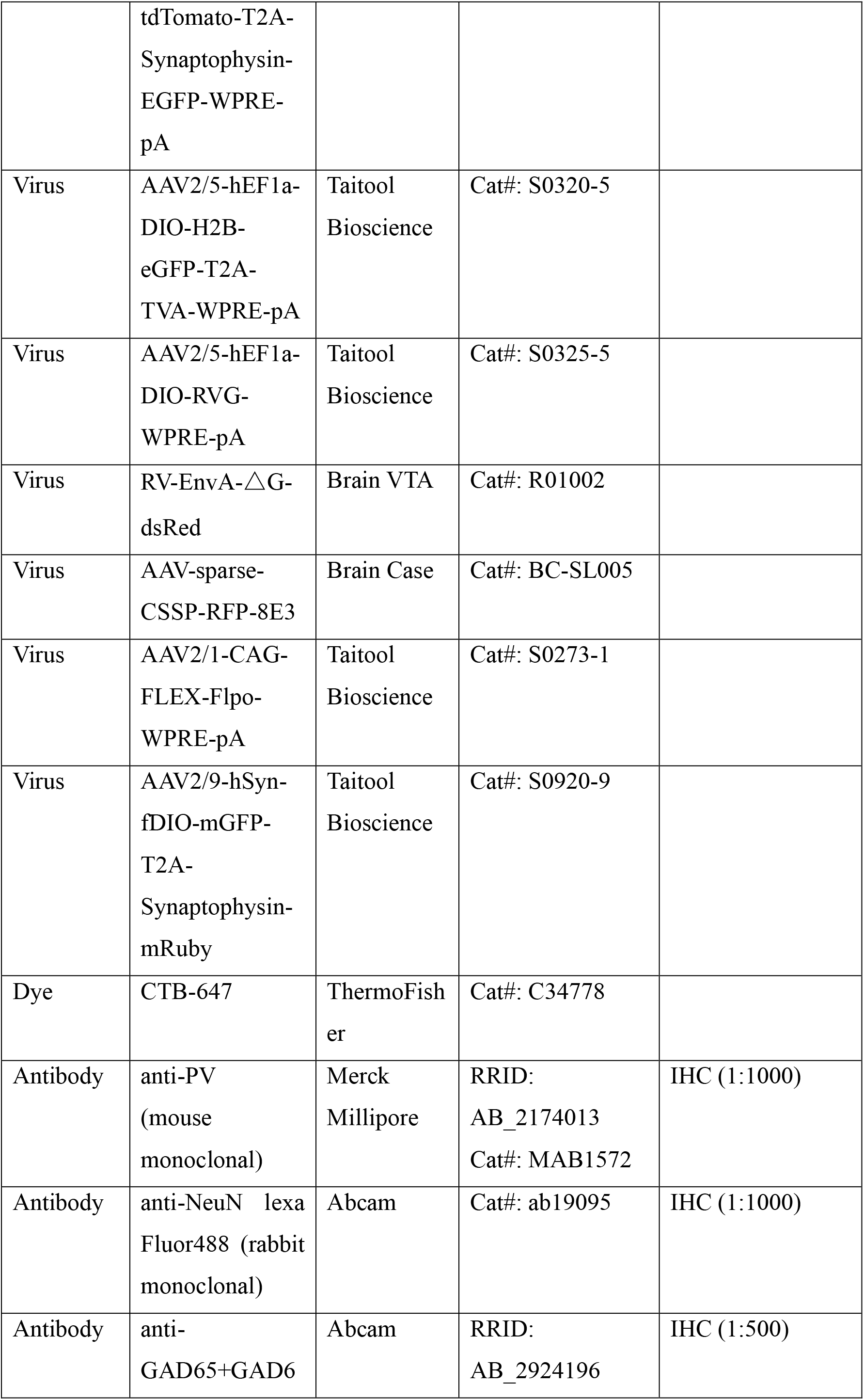
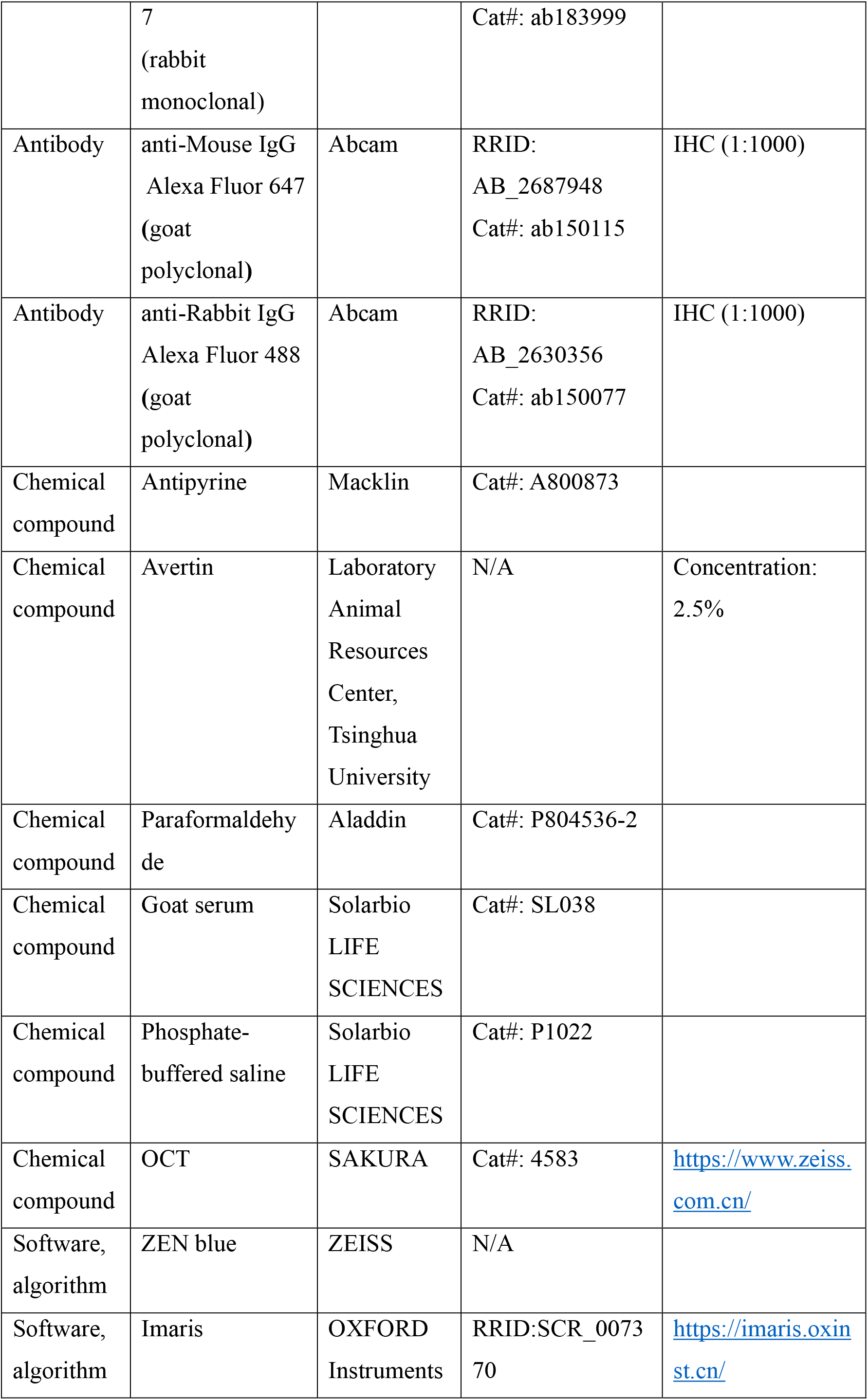
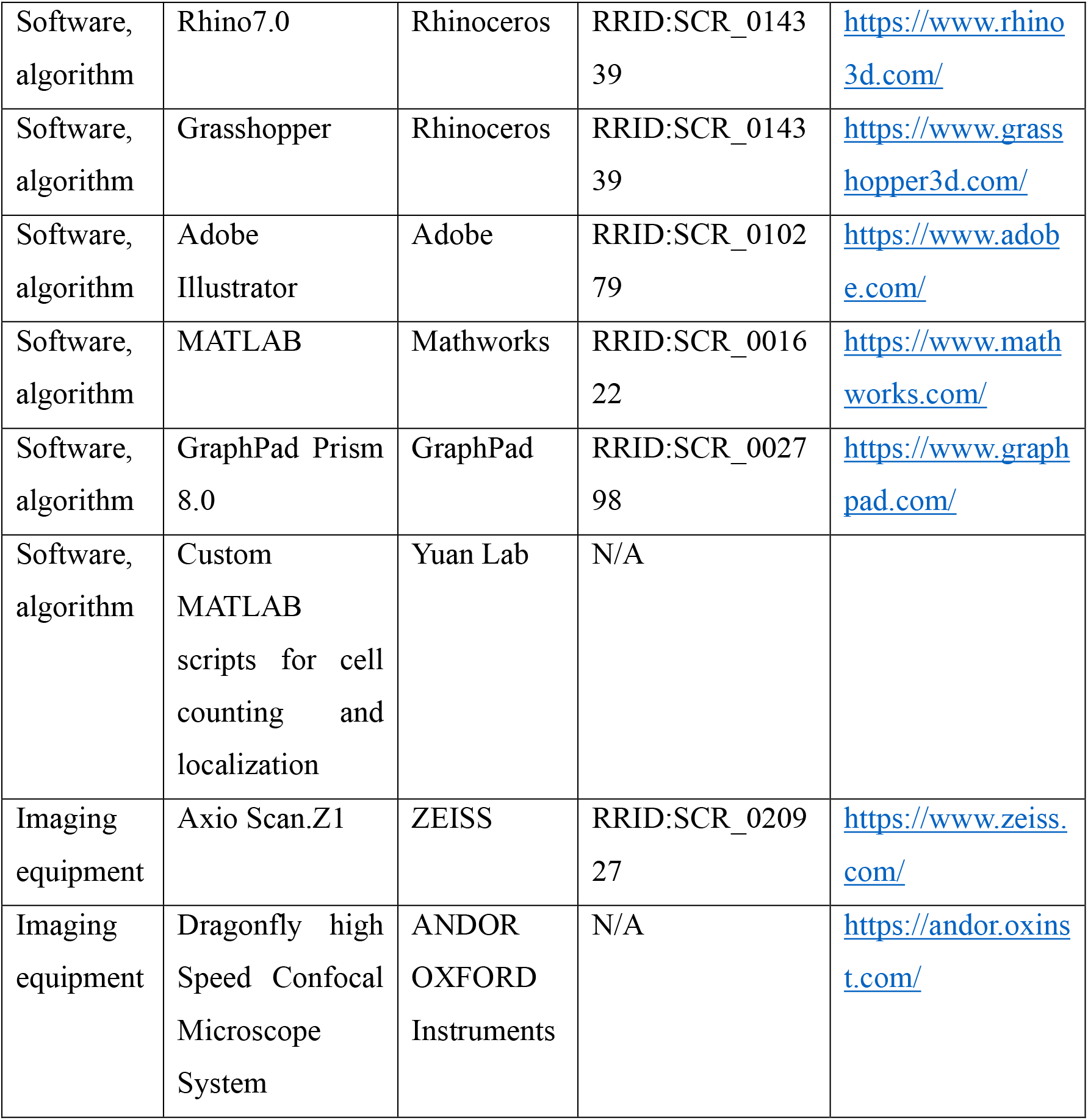
Key resources table.

### EXPERIMENTAL DETAILS

#### Abbreviations for brain regions

The majority of abbreviations for brain regions used in this study were from the Allen Mouse(https://scalablebrainatlas.incf.org/main/coronal3d.php?template=ABA_v3#downloads) and the Mouse Brain in Stereotaxic Coordinates, second edition by Franklin, K. B. J. and Paxinos, G.

| Abbreviation | Definition |
| --- | --- |
| AAV | Adeno-associated virus |
| AUD | Auditory cortical area |
| AUDd | Dorsal auditory cortical area |
| AUDp | Primary auditory cortical area |
| AUDv | Ventral auditory cortical area |
| CN | Cochlear nuclei |
| CO | Cytochrome oxidase |
| CUN | Cuneiform nucleus |
| DCN | Dorsal cochlear nucleus |
| FS | Fast spiking |
| GAD | Glutamic acid decarboxylase |
| GPe | External segment of the globus pallidus |
| IC | Inferior colliculus |
| ICC | Central nucleus of the inferior colliculus |
| ICD | Dorsal cortex of the inferior colliculus |
| ICE | External cortex of the inferior colliculus |
| KDE | Kernel density estimation |
| LD | Lateral dorsal nucleus of thalamus |
| MGB | The medial geniculate body |
| MGBv | Ventral division of the medial geniculate body |
| MGBd | Dorsal division of the medial geniculate body |
| MGBm | Medial division of the medial geniculate body |
| MRN | Midbrain reticular nucleus |
| NLL | Nucleus of the lateral lemniscus |
| NSTR | Neurotensin receptor |
| OCT | Optimal cutting temperature |
| PAG | Periaqueductal gray |
| PB | Parabrachial nucleus |
| PBGN | Parabigeminal nucleus |
| PBS | Phosphate-buffered saline |
| PFA | Paraformaldehyde |
| POL | Posterior limiting nucleus of the thalamus |
| PPN | Pedunculopontine nucleus |
| PIN | Posterior intralaminar thalamic nucleus |
| PV | Parvalbumin |
| RI | Rapid inactivating |
| RS | Regular spiking |
| RV | Rabies virus |
| SC | Superior colliculus |
| SCm | Motor layers of the superior colliculus |
| SCig | Intermediate gray layer of the superior colliculus |
| SNr | Substantia nigra, reticular part |
| SOC | Superior olivary complex |
| SOCm | Medial part of the superior olivary complex |
| SOCl | Lateral part of the superior olivary complex |
| SOM | Somatostatin |
| SPF | Subparafascicular nucleus |
| SSs | Supplemental somatosensory cortical area |
| TEa5 | Temporal association areas, layer 5 |
| TS | Tail of the striatum |
| VIP | Vasoactive intestinal peptide |
| WT | Wild type |
| ZI | Zona incerta |

#### Animals

All animal care procedures and experiments described in this study were conducted in strict accordance with the ethical guidelines and regulations set forth by the Institutional Animal Care and Use Committee (IACUC) at Tsinghua University, Beijing, China. Prior to and following surgical procedures, all mice were maintained under standard environmental conditions of temperature, humidity, and light/dark cycles at the Laboratory Animal Resources Center, Tsinghua University. Appropriate measures were taken to ensure animal welfare and minimize potential discomfort or harm. Adult mice (∼2 months, Background strain C57BL/6J) were used for tracing and immunofluorescence staining studies, while juvenile mice (3-8 weeks, Background strain C57BL/6J) of both sexes were used for whole-cell patch experiments. C57BL/6J (wild-type, WT) mice were procured from Wei Tong Li Hua Experimental Animal Co., Ltd (Beijing, China), and all transgenic mice were obtained from the Jackson Laboratory. The PV-IRES-Cre mice (B6.129P2-Pavlb^tm1(cre)Arbr^/J, Jackson Laboratory, Stock#: 017320) and SOM-IRES-Cre mice (Sst^tm2.1(cre)Zjh^/J, Jackson Laboratory, Stock#: 013044) (*Taniguchi et al., 2011*) were utilized for labeling PV+ and SOM+ neurons in the IC, respectively. The Vgat-Cre mice (Slc32a1^tm2(cre)Lowl^/J, Jackson Laboratory, Stock#: 016962) (*Vong et al., 2011*) were used for labeling inhibitory neurons in the IC. In addition, the Ai14 mice (B6. Cg-Gt (ROSA)26Sor^tm14(CAG-tdTomato)^ ^Hze^/J, Jackson Laboratory, Stock#: 007914) (*Madisen et al., 2010*), also known as Cre-reporter mouse line, were crossed with SOM-Cre (SOM× Ai14), PV-Cre (PV×Ai14) and Vgat-Cre (Vgat×Ai14) mice to achieve cell-type specific expression of red fluorescent protein in neurons within the IC in this study.

#### Stereotaxic surgeries

The mice were first anesthetized with 2.5% avertin (300 mg/kg, i.p.) and placed in a stereotaxic apparatus (RWD, Shenzhen, China). To ensure the comfort of the mice, erythromycin eye ointment was administered to prevent eye dryness, and an electric heating pad was used to maintain their body temperature during the surgical process. The skin over the skull midline was then carefully incised with sterilized scissors, and forceps were used to expose the bregma, lambda, and skull surface. Next, a small hole was performed above the IC or auditory thalamus, and a micro-syringe pump with a glass pipette (WPI, USA) was used to slowly inject viruses (35nl/min) into the IC or auditory thalamus. The following coordinates (in mm) were used (Paxinos and Franklin, 1997): -3.00 AP, -2.10 ML, -3.00 DV for the MGBv; -3.00 AP, -1.75 ML, -2.80 DV for the POL; -1.20 AP1, -0.60 ML, -0.70 DV for the ICD; -1.20 AP1, -1.00 ML, -0.8 DV for the ICC; -1.20 AP1, -1.60 ML, -0.70 DV for the ICE (AP is relative to bregma for auditory thalamus nuclei, AP1 is relative to lambda for the IC; ML is relative to the midline; DV is relative to the brain tissue surface above the targeted brain areas). The glass pipette remained in place for an additional 5 minutes after injection before being slowly withdrawn. The skin was sutured after retracting the pipette. After surgery, the mice were allowed to recover on a heated pad until they were ambulatory, and then they were returned to their home cage. The mice underwent a post-surgery recovery period of approximately 2-3 weeks to allow for adeno-associated virus (AAV) virus infection and gene expression.

#### Virus injection and tracing

For cell type-specific anterograde tracing, we employed an injection of 150nl AAV2/9- hSyn-Flex-Synaptophysin-mRuby (Cat# WY3330, 3.6E+12V.G/ml, Taitool Bioscience, Shanghai, China) into the IC of PV-IRES-Cre or SOM-IRES-Cre mice to trace the projection of PV+ or SOM+ neurons in all subdivisions of the IC. Furthermore, we applied the same virus (30-50nl) injection as local as possible in individual IC subdivision to investigate how PV+ and SOM+ neurons in each subdivision of the IC participate in the tectothalamic pathways. In addition, we conducted control experiments in WT mice by performing AAV2/9-hSyn-Flex-Synaptophysin-mRuby injection mixed with CTB-647 (C34778, ThermoFisher) into the IC, aiming to confirm the crucial reliance of mRuby expression on Cre (*supplement figure 2C*).

For trans-synaptic anterograde tracing, we injected 80-100nl scAAV-hSyn-Cre (Cat# S0292-1, 1.25E+13 V.G/ml, Taitool Bioscience, Shanghai, China) into the IC of the WT mice. At the same time, we preformed localized injection of AAV2/5-Flex-tdTomoto-T2A-Synaptophysin-EGFP (Cat# S0161-5, 5.3+12E V.G/ml, Taitool Bioscience, Shanghai, China) into the MGBv or AAV2/9-Flex-Synaptophysin-mRuby into the POL, to determine the downstream target of IC-MGBv or IC-POL pathways.

To identify the input of PV+ or SOM+ neurons in the IC, we adopted a widely used rabies virus (RV)-mediated retrograde monosynaptic tracing strategy. Briefly, mixed Cre-dependent AAV helper viruses AAV2/5-hEF1a-DIO-H2B-eGFP-T2A-TVA-WPRE-pA (Cat#S0320-5, 7.3+12E V.G/ml, Taitool Bioscience, Shanghai, China) and AAV2/5-hEF1a-DIO-RVG-WPRE-pA (Cat#S0325-5, 5.5+12E V.G/ml, Taitool Bioscience, Shanghai, China) were injected into the IC of PV-IRES-Cre or SOM-IRES-Cre mice to selectively express TVA and RG in PV+ or SOM+ neurons (*Figure 3A,* Day 1). Three weeks later, 100nl of pseudotyped RV equipped with the TVA selective avian ASLV type A (EnvA) (RV-EnvA-△G-dsRed; Cat#R01002, 2.00+08E IFU/ml, BrainVTA, Wuhan, China) was injected into the same injection sites (*Figure 3A,* Day 21) to enable dsRed expression in the input neurons of PV+ or SOM+ starter neurons (*Figure 3B,* PV+; *Figure 3C*, SOM+; AAV-Helper, green; Input neurons, red; Starter neurons, yellow). For control experiments, we injected the same set of viruses with the same procedures into the IC of SOM-IRES-Cre mice without including either AAV-DIO-RG (*supplement figure 3A,* N=4 mice) or AAV-DIO-EGFP-TVA (*supplement figure 3B,* N=4 mice), and no input neurons were observed in the control mice, validating the dependency of retrograde tracing on RG or TVA. The mice were sacrificed after RV expression and retrograde spreading for another 7 days (*Figure 3A,* Day 28).

To visualize the morphology of IC^SOM+^ and IC^PV+^ neurons, we performed injections of AAV-sparse-CSSP-RFP-8E3 (Cat#BC-SL005, 5+12E V.G/ml, Brain Case, Shenzhen, China) into the IC of SOM-IRES-Cre and PV-IRES-Cre mice, respectively. We also used a combination of AAV2/1-CAG-FLEX-Flpo-WPRE-pA (Cat#S0273-1, 2.73+10E V.G/ml, Taitool Bioscience, Shanghai, China) and AAV2/9-hSyn-fDIO-mGFP-T2A-Synaptophysin-mRuby (Cat#S0920-9, 6.8+12E V.G/ml, Taitool Bioscience, Shanghai, China) for injections into the IC of SOM-IRES-Cre or PV-IRES-Cre mice to visualize the morphology of IC^SOM+^→POL or IC^PV+^→MGBv axon terminals.

#### Histology and Immunochemistry

Animals were administered an overdose of 2.5% avertin (350mg/kg, i.p.) and underwent transcardially perfused with 0.9% saline, followed by 4% paraformaldehyde (PFA, P804536-2, Aladdin) in 0.01M phosphate-buffered saline (1×PBS, Solarbio LIFE SCIENCES). The brains were then extracted, post-fixed in 4% PFA at 4℃ overnight, and subsequently immersed in 30% sucrose for dehydration and cryoprotection until they sank. The brains were frozen in optimal cutting temperature compound (OCT, Sakura Tokyo, Japan), and coronal slices were obtained using a freezing microtome (CM1950, Leica Biosystems, Germany) at a thickness of 50μm, covering the whole brain. For brain-wide cell counting, every other slice was collected and mounted on gelatin-coated slides. For antibody immunofluorescent staining, coronal brain sections of 40-50μm were collected into multiple well plates (Corning® Costar®) coated with an ultra-low attachment material and filled with 1×PBS. The collected brain sections were rinsed three times for 5 minutes (3×5min) each with 1×PBS buffer on a shaker. Next, brain sections were incubated for 3×15min with 0.3% Triton 100 in 1×PBS for permeabilization, followed by a 60min incubation with 5% goat serum (SL038, Solarbio) to block unspecific binding of the antibodies. The brain sections were then incubated with primary antibodies overnight at 4℃, followed by 3×15min washes with 1×PBS, and incubated with secondary antibodies for 2 hours. The sections were then thoroughly washed with 1×PBS and cover-slipped with 50% glycerol mounting medium. We used the primary antibody of anti-PV mouse monoclonal antibody (1:1000, MAB1572, Merck Millipore) combined with the secondary antibody of anti-Mouse polyclonal goat antibody conjugated with Alexa Fluor 647 (1:1000, ab150115, Abcam) to label PV+ neurons in the IC. We also employed the primary antibody of GAD65+GAD67 rabbit monoclonal antibody (1:500, ab183999, Abcam) with anti-Rabbit polyclonal goat secondary antibody conjugated with Alexa Fluor 488 (1:1000, ab150077, Abcam) to label GAD+ neurons in the IC. We used anti-NeuN rabbit monoclonal antibody conjugated with Alexa Fluor488 (1:1000, ab19095, Abcam), which was used without secondary antibody incubation, to label NeuN+ neurons in the IC. To validate the specificity of each primary antibody, we performed control experiments following the same procedures as described previously, except that 1×PBS was substituted for the primary antibodies.

#### Acute brain electrophysiology and recordings

SOM× Ai14 and PV×Ai14 mice (3-8 weeks old, both sexes) were deeply anesthetized by intraperitoneal administration of 2.5% avertin (300mg/kg) and were then perfused through the heart using NMDG-HEPES (*Ting et al., 2018*) artificial cerebrospinal fluid (ACSF) consisting of (in mM): 92 NMDG, 2.5 KCl, 1.25 NaH_2_PO_4_, 30 NaHCO_3_, 20 HEPES, 25 Glucose, 2 Thiourea, 5 Na-ascorbate, 3 Na-pyruvate, 0.5 CaCl_2_·2H_2_O, and 10 MgSO_4_·7H_2_O. The pH of the solution was adjusted to 7.3-7.4 using 5M hydrochloric acid, and the osmolality was 300-310 mOsmoles/kg. The solution was pre-chilled at 4°C and bubbled with carbogen (95% O_2_ and 5% CO_2_) beforehand. We obtained approximately four coronal slices, each with a thickness of 300 µm, using a vibratome (Leica VT1200S, Germany). Subsequently, the slices were transferred to a pre-warmed (34°C) beaker filled with NMDG-HEPES ACSF for a protective recovery spike-in procedure. Afterwards, these slices were transferred into HEPES-holding ACSF consisting of (in mM): 92 NaCl, 2.5 KCl, 1.25 NaH_2_PO_4_, 30 NaHCO_3_, 20 HEPES, 25 Glucose, 2 Thiourea, 5 Na-ascorbate, 3 Na-pyruvate, 2 CaCl_2_·2H_2_O, and 2 MgSO_4_·7H_2_O, with the pH titrated to 7.3-7.4 using 10M NaOH. The holding chamber was equilibrated at room temperature for a minimum of one hour prior to the recording session and was maintained at this temperature throughout the one-day experiment. Individual slices were transferred to the recording chamber as required. Slices were transferred to a submerged recording chamber on the stage of an upright microscope (Olympus BX51WI, Japan). The chamber was continually perfused (2-3ml/min) with recording ACSF solution under constant oxygen and containing (in mM): 124 NaCl, 2.5 KCl, 1.25 NaH_2_PO_4_, 24 NaHCO_3_, 12.5 Glucose, 5 HEPES, 2 CaCl_2_·2H_2_O, and 2 MgSO_4_·7H_2_O. Individual SOM+ or PV+ neurons were identified under differential interference contrast (DIC) or fluorescence and a 40× water immersion objective with an infrared CCD camera (DAGE MTI IR-1000E, USA). Whole-cell recordings were obtained using 8-12 MΩ pipettes pulled from borosilicate glass by a puller (P97, Sutter Instruments), with an in-pipette solution containing 122.5 mM K-glucose, 12.5 mM KCl, 10 mM HEPES, 2 mM Na_2_ATP, 0.3 mM Na_2_GTP, 2 mM MgCl_2_, and 8 mM NaCl. Electrophysiological signals were amplified using a MultiClamp700B amplifier (Molecular Devices, USA), digitized (Axon Digidata1440A, USA) at 20-50 kHz, and filtered at 2-10 kHz.

#### Rapid and non-scaling on-slide tissue clearing

In this study, we developed an innovative and efficient on-slide tissue clearing methodology that enables rapid and non-scaling processing for visualization of tectothalamic terminal structures. To achieve high-throughput and high-precision imaging, we combined this technique with high-speed spinning-disk confocal imaging. Following prefusion using 4% PFA, the mouse brain was placed in a balanced cryopreserved solution containing 10% Dimethyl sulfoxide, 3% Sucrose, and 1×PBS solution for two days. The OCT-embedded brains were then cryosectioned at approximately -25℃. The resulting brain slices were placed onto gelatin-coated slides and incubated in a high-humidity environment at 37°C for two hours. After thorough rinsing with 1×PBS solution was conducted for 10 minutes, the brain slices were subjected to tissue clearing by treating them with a 55% antipyrine (A800873, Macklin) and 0.1% DAPI (D9542, Sigma) on-slide clearing mounting solution dissolved in deionized water for 2-3 minutes. The same clearing solution was used for cover-slipping the brain slices. Prior to imaging, the slides were stored in a dark environment at room temperature. This optimized protocol effectively preserved tissue morphology and ensured the acquisition of reliable imaging data for our analysis (see *supplement figure 4C*).

#### Imaging and fluorescent signal recognition

To enable brain-wide cell counting and analysis of neuron distribution, the slides were imaged using the Axio Scan.Z1 (Zeiss Axio Scan, Germany) equipped with a 10× objective, allowing for efficient scanning of large areas and capturing an overview of the samples. For higher-resolution imaging and precise visualization of microscale structures and immunofluorescence staining in brain slices, we utilized the high-speed confocal microscopy system (ANDOR, OXFORD Instruments, Britain) with a 20×objective. This enabled us to obtain detailed and accurate information about the cellular colocalization and fine structural features. To further investigate tectothalamic terminal structures, we employed a 40× objective with a 0.5μm z-step, enabling us to capture high-resolution images of synapses. During confocal imaging, we ensured a single-field resolution of 2048×2048, acquired images with an average of two scans, and used 16-bit imaging to preserve the full dynamic range of the data. Finally, to generate complete three-dimensional images, automatic stitching was employed to seamlessly merge multiple fields of view, resulting in comprehensive and detailed visualization of the samples.

By employing a combination of automated identification and manual assessment, we identified fluorescence signals corresponding to neurons and synaptic structures. To achieve this, we utilized the Spots tool in Imaris9.0 software (OXFORD Instruments, https://imaris.oxinst.cn/). Size thresholds were set for the two distinct types of fluorescent signals, with a range of 0.5μm to 2μm for synaptic structures and 10μm to 12μm for neurons. Manual intervention was applied to supplement or remove misidentified or missed fluorescent signals. In particular, manual adjustments were necessary when the fluorescent intensity of proximal dendrites or axons was exceptionally high, making it challenging for the software to accurately identify the neuron soma or distinguish neighboring neurons. Subsequently, the positions of the fluorescent markers on the corresponding images, as well as the images themselves, were imported into Rhinoceros7 software (Rhinoceros, https://www.rhino3d.com/) and aligned with a widely utilized standard brain atlas obtained from the Allen Reference Atlas(AllenInstitute,https://scalablebrainatlas.incf.org/main/coronal3d.php?template=ABA_v3#). Statistical analysis was performed using custom MATLAB scripts (Yuan Lab, MathWorks) for cell counting and localization, as well as custom Grasshopper scripts (Yuan Lab, Rhinoceros) for the quantification of auditory thalamic terminals, to conduct further analysis.

## Data analysis

### Kernel density estimation

The density distribution of IC neurons was analyzed using kernel density estimation (KDE) with an adjusted quad kernel densitometry method (*BW, 1986*). This approach allowed us to estimate the density distribution of dot signals at any given location (x, y) within the identified neurons. The calculation was derived from the following formula.

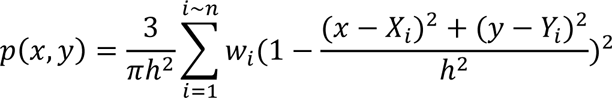

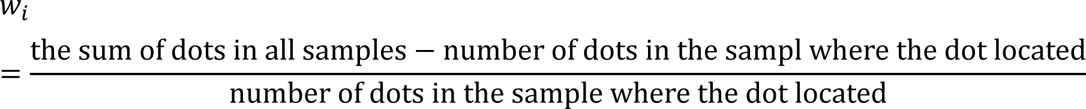

The bandwidth (h) in the KDE was determined using the following rule, where Dm represented the median value of the distance between the dots and their average center.

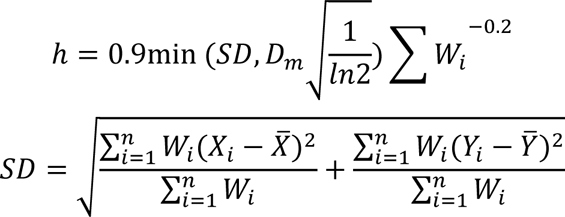

The density distribution heatmap was generated by assigning the values of p (x, y) to corresponding color levels.

### Quantification estimation of tectothalamic synaptic terminals

To quantitatively analyze the tectothalamic pathways, we focused on the brain slice at the AP=-2.97mm coordinate, which represents a typical section of the auditory thalamus. By using “fluorescent signal recognition” method described above, we identified the axon terminals in different subdivisions of the auditory thalamus and determined the total number of terminals originating from IC^SOM+^ or IC^PV+^ projection neurons. We then calculated the percentage (P) and weighted density (D) of synaptic terminals in each auditory thalamus subdivision to assess the strength of the connection between IC neurons and each auditory thalamus nucleus. Specifically, P represents the percentage of terminals in a given auditory thalamus subdivision relative to the total number of terminals across all subdivisions. We only presented the P values for IC^SOM+^ and IC^PV+^ neuron projections in the MGBv, MGBd, POL, and PIN, as their projections to other subdivisions of the auditory thalamus are minimal (*Figure 3, D-I, 3^rd^ column, bar charts*). Additionally, we included the weighted density (D) due to the significant differences in the size (A) of these four auditory thalamus subdivisions (*Figure 3, D-I, 3^rd^ column, line graphs*).

To visualize the distribution of IC^SOM+^ and IC^PV+^ terminal projections to the auditory thalamus, we conducted a random and even sampling of 3000 terminals from each mouse brain slice at the -2.97mm coordinate. These selected terminals were used as representative samples to illustrate the average distribution pattern across the entire sample (*Figure 3, D-F, G, I, 2^nd^ column*). We observed a low number of terminals originating from SOM+ neurons in the ICC projecting to the thalamus, and this dataset illustrated the distribution of 300 terminals representing the projection of ICC^SOM+^ neurons to the auditory thalamus (*Figure 3*, H, *2^nd^ column)*

### Data analysis for retrograde tracing

To evaluate the similarity in the number of RV-mediated retrograde monosynaptic tracing labeled neurons among input sources, we used Pearson’s correlation coefficient of normalized inputs, calculated as the total number of input neurons from a specific brain region divided by the total number of input neurons.

### Electrophysiology data analysis

All electrophysiological data were analyzed using our custom-written MATLAB program. Traces obtained from PV+ or SOM+ neurons, depicting their responses to current injections, were utilized to extract several electrophysiological parameters. The resting membrane potential was determined as the mean value from trials without any injected current (*Figure 5C*). The rheobase was defined as the minimum current required to elicit the first single action potential in a neuron (*Figure 5D*). The time constant was derived from a single exponential decay function fitted between 10% and 95% of the response to a -80pA current injection (*Figure 5F*). Sag was quantified by measuring the voltage response during a hyperpolarizing current injection of -80pA (*Figure 5E*).

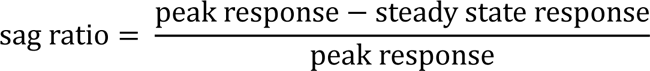

To investigate the electrophysiological characteristics of PV and SOM neurons and to present them in an unbiased manner, we utilized t-Distributed Stochastic Neighbor Embedding (t-SNE) with the MATLAB implementation. This technique allowed us to project the relevant electrophysiological parameters onto a two-dimensional space. In order to obtain optimal results, we carefully adjusted the hyperparameters considering the size of our dataset (PV: 28, SOM: 22). Specifically, we set the perplexity to 10, the learning rate to 1000, and employed the cosine distance metric for the t-SNE analyses (*Figure 5G*).

### Terminal size analysis

We utilized synaptophysin-mRbuy labeling to identify tectothalamic axon terminals in the auditory thalamus. By combining the turntable confocal microscope with our previously described “Rapid and non-scaling on-slide tissue clearing” method (see detailed methods above), we successfully achieved high-throughput three-dimensional fluorescence imaging at the single-synapse scale in large-scale brain regions. Our data processing workflow involved several steps. First, we performed deconvolution based on the point spread function of the microscope system. Next, background subtraction was applied, and the volume of fluorescent signals was reconstructed using a fluorescence intensity threshold. If necessary, point recognition algorithms were used to separate closely adhered signals and generate models for each fluorescent dot signal. Finally, we extracted the fluorescence intensity and volume information from each three-dimensional reconstructed object.

To accurately quantify the size of axon terminals using a fluorescence microscope, we employed fluorescent microbeads with known sizes (e.g., 0.5μm, 1μm, 2μm, and 3μm) that closely resembled the size range of real synaptic terminals (*De Banne et al., 2011*). By imaging these microbead samples, which exhibited varying fluorescence intensities achieved through gradient fluorescence quenching, we established a relationship function between the parameters of fluorescent signals obtained from the fluorescence microscope and the corresponding real sizes of the microbeads. The microbeads were injected into either agarose gel or the mouse brain, and their reconstructed volumes were determined, enabling us to establish a robust correlation between the fluorescence signal parameters and the actual sizes of synaptic terminals. Subsequently, we performed a linear fitting analysis by taking the negative reciprocal of the mean fluorescence intensity of the microbeads and their corresponding reconstructed volumes. The results revealed a strong positive linear correlation between these two parameters. Based on this correlation, we constructed a surface representation in three-dimensional space by lofting a fitting line, which accurately described the relationship between the real size of the targeted microbeads and their reconstructed volumes. This surface provided a comprehensive depiction of the distribution of both the negative reciprocal of the fluorescence intensity and the reconstructed size, allowing for precise estimation of the actual sizes of synaptic terminals based on their fluorescence signal characteristics. The established model was then applied to reconstruct the volumes of all fluorescent microbeads of different sizes (0.5μm, 1μm, 2μm, and 3μm). As anticipated, we observed three distinct peaks in the diameter distribution function after identifying the peak values using Fourier transform and fitting Gaussian distributions to the reference peaks. Notably, the distribution of reconstructed volumes exhibited a clear superposition of three normal distributions, with each mean value corresponding to the true size of the microbeads. We further calculated the mean values and areas under the curve of the fitting normal distributions. The results demonstrated that the areas under the fitting distribution curves (or peaks) accurately reflected the real proportions of the three given microbead sizes. This confirms that our methodology effectively represents the true size and number of fluorescent microbeads. In conclusion, our approach not only accurately estimates the real size of fluorescent signals ranging from 0.5μm to 2μm but also accurately captures the distribution of fluorescent signals with mixed sizes.

### Quantification and statistical analysis

Statistical analysis and tests were conducted using GraphPad Prism (GraphPad Software, USA) or MATLAB (MathWorks, USA) with two-tailed. The normality of data was assessed using the Shapiro-Wilk test. For normally distributed data, a t-test was employed, while non-parametric Mann-Whitney test was used for non-normally distributed data. In cases of multiple comparisons, post-hoc tests such as Bonferroni and Dunn’s tests were utilized for One-way ANOVA and Kruskal-Wallis H-test, respectively. The data are presented as means±SEMs in both figures and text. Paired tests for two groups were performed using the paired t-test for normally distributed data and the Wilcoxon matched-pairs signed rank test for non-normally distributed data. Multiple paired tests were analyzed using two-way ANOVA with Geisser-Greenhouse correction. The statistical details, including the specific tests used, statistics, significance levels, sample sizes, animal numbers are provided in the figure legends. Significance levels are denoted as *P<0.05, **P<0.01, ***P<0.001, ****P<0.0001, and n.s. for non-significant (P>0.5).

---

**supplement Figure 1.**
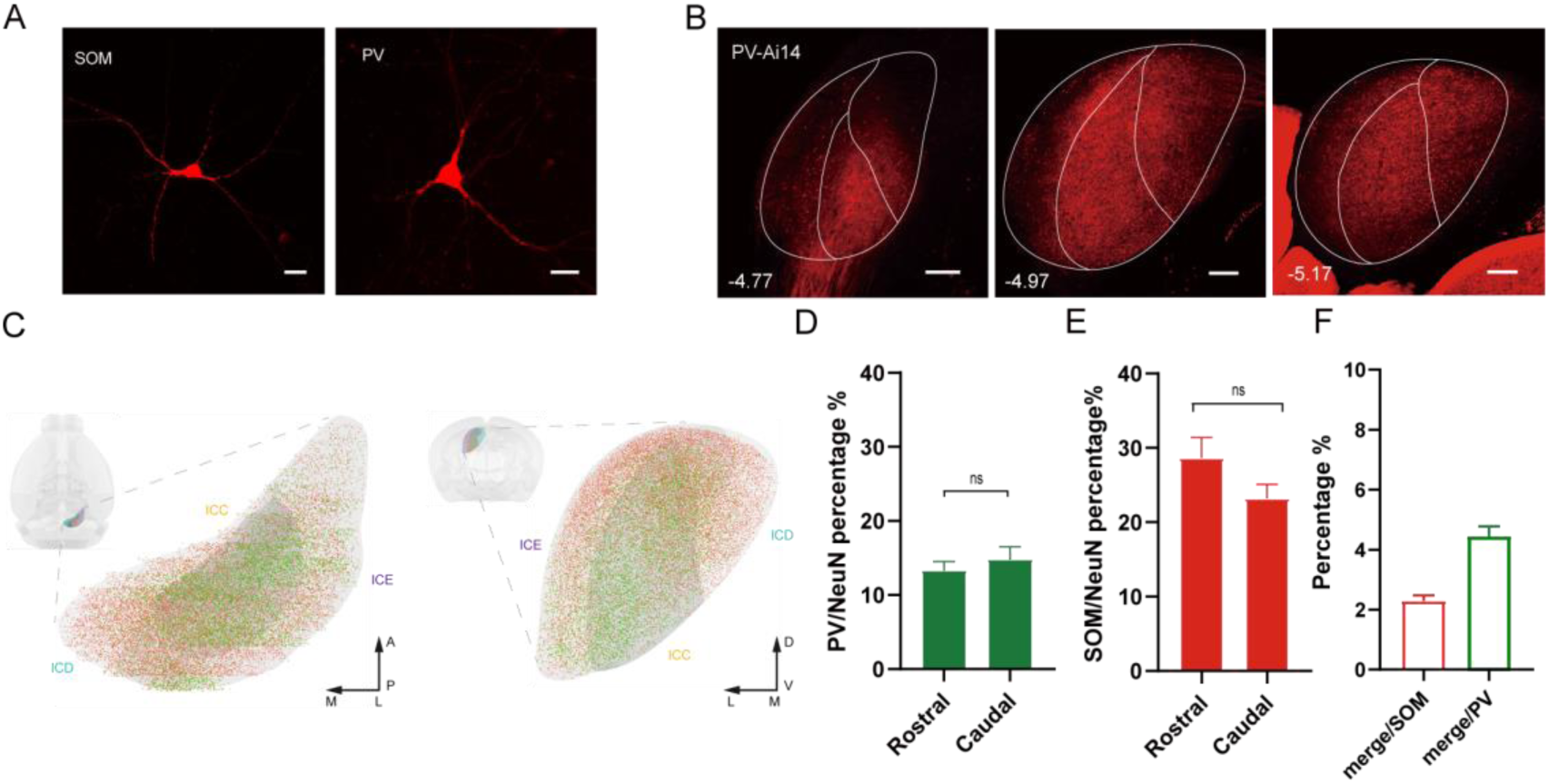
Related to Figure 1. The morphology and distribution of PV+ and SOM+ neurons in the IC. (A) Confocal 10× images showing the morphology of IC^SOM+^ and IC^PV+^ neurons by sparse labeling. Scale bars, 50μm. (B) Example images showing dense PV+ neurites in the ICC in PV-IRES-Cre×Ai14 mice slices. Scale bars, 200μm. (C) Illustrative diagrams showing the distribution of PV+ neurons (green dots) and SOM+ neurons (red dots) in different directions within the IC. AP, anterior-posterior; ML, medial-lateral; DV, dorsal-ventral. (D) Percentage of PV+ neurons of NeuN neurons in rostro-caudal axis of the IC. n=8 slices from N=4 mice, unpaired t-test, p>0.5. (E) Percentage of SOM+ neurons of NeuN neurons in rostro-caudal axis of the IC. n=8 slices from N=4 mice, unpaired t-test, p>0.5. (F) Percentage of PV+ and SOM+ co-expression neurons (merge) among PV+ or SOM+ counted neurons; n=6 slices from N=2 mice. Data are means± S.E.M.

**supplement Figure 2.**
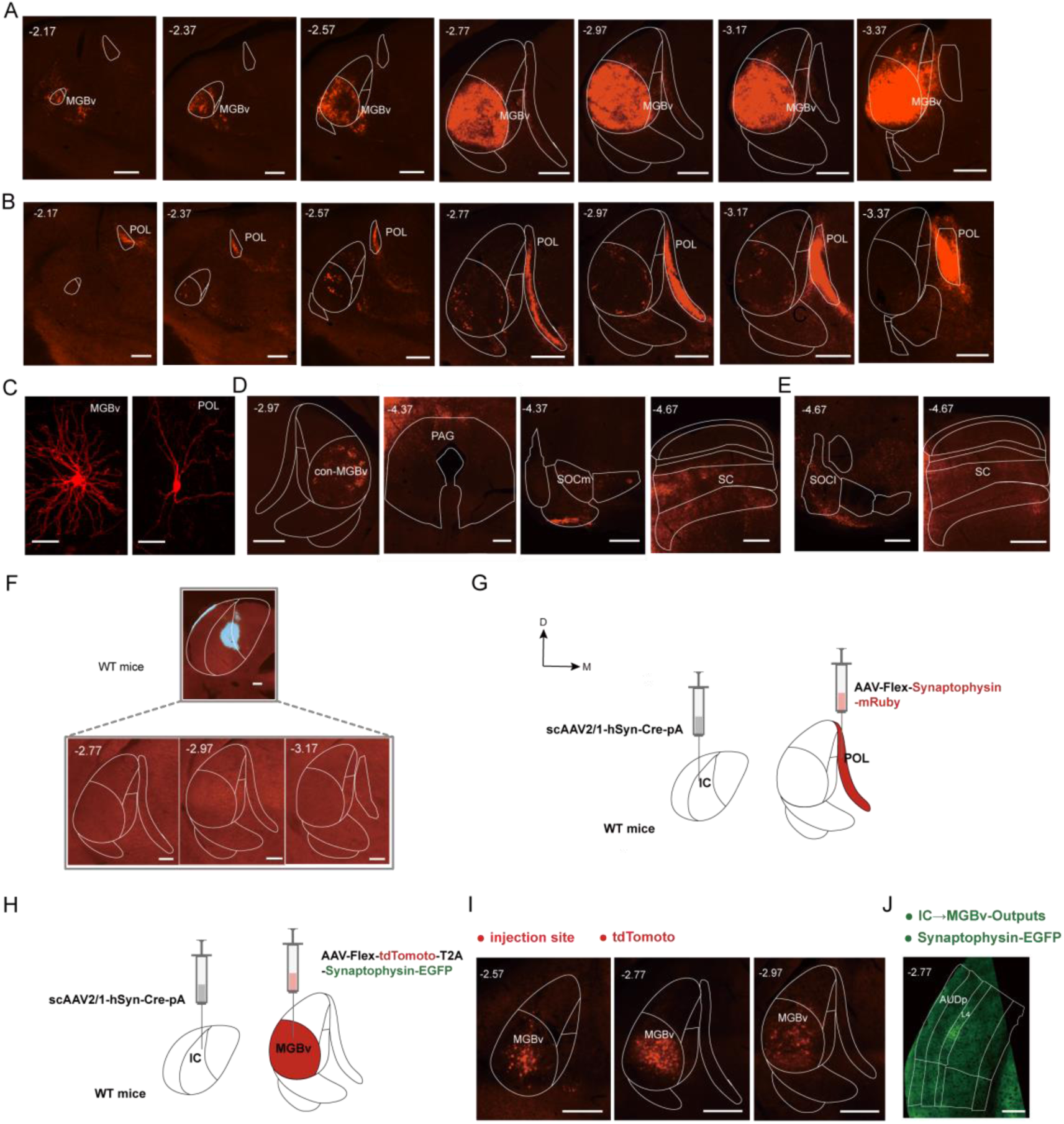
Expanded data to Figure 2. Projections of PV+ and SOM+ neurons in the IC. (A) Example images showing when the virus was injected in a subdivision-independent manner into the IC of PV-IREW-Cre, the projections to the ipsilateral auditory thalamus from anterior to posterior coordinate. Scale bars, 300μm. (B) Example images showing when the virus was injected in a subdivision-independent manner into the IC of SOM-IRES-Cre, the projections to the ipsilateral auditory thalamus from anterior to posterior coordinate. Scale bars, 300μm. (C) Confocal 10× images showing the morphology of MGBv and POL neurons by sparse labeling. Scale bars, 50μm. (D) Example images showing when the virus was injected in a subdivision-independent manner into the IC of PV-IRES-Cre, the projections to brain regions other than the ipsilateral auditory thalamus. Distribution of axonal terminals in the contralateral MGBv (con-MGBv), PAG, SOCm and SC. Scale bars, 300μm. (E) Example images showing when the virus was injected in a subdivision-independent manner into the IC of SOM-IRES-Cre, the projections to brain regions other than the ipsilateral auditory thalamus. Distribution of axonal terminals in the SOCl and SC. Scale bars, 300μm. (F) Validation of the dependency of mRuby expression on Cre. Upper panel, example image showing injection AAV-Flex-synaptophysin-mRuby with CTB-647 into the IC of WT mice. Scale bar, 200μm. Lower panel, the projections to the auditory thalamus. Scale bars, 300μm. (G) Experimental procedures for anterograde tracing of IC→POL pathway. Left panel, scAAV2/1-Cre-PA virus was injected into the IC of WT mice. Right panel, targeted injections of AAV2/9-Flex-Synaptophysin-mRuby into the POL. D, dorsal; M, medial. (H) Experimental procedures for anterograde tracing of IC→MGBv pathway. Left panel, scAAV2/1-Cre-PA virus was injected into the IC of WT mice. Right panel, targeted injections of AAV2/5-Flex-tdTomoto-T2A-Synaptophysin-EGFP into the MGBv. (I) Example images showing expression of mRuby localized in the MGBv neurons. Scale bars, 300μm. (J) Example images showing major target brain region of the synaptic terminals of MGBv neurons receiving IC inputs. Scale bar, 300μm.

**supplement Figure 3.**
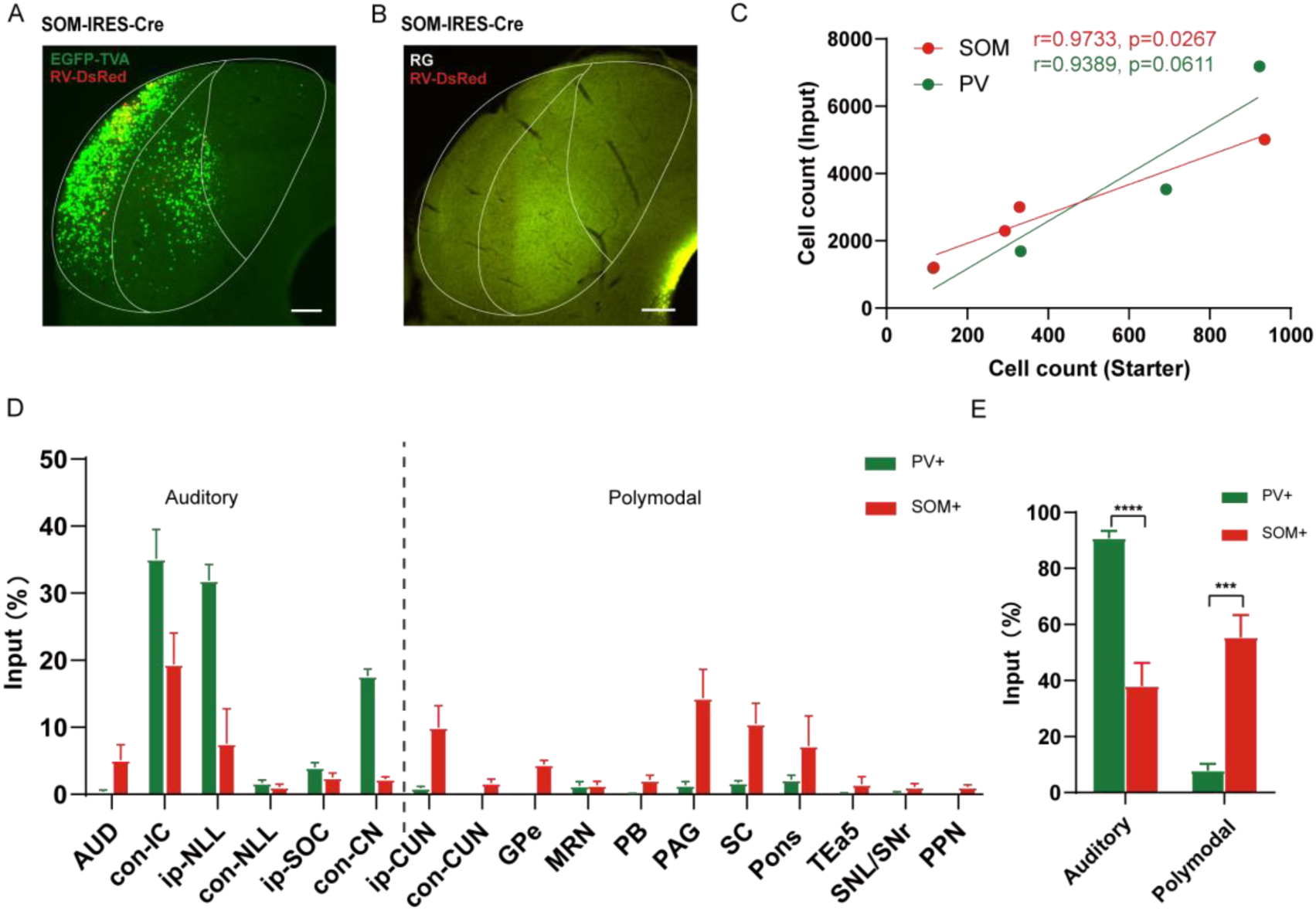
Expanded data to Figure 3. Inputs of PV+ and SOM+ neurons in the IC. (A) Validation of RG-dependence of RV-mediated monosynaptic retrograde tracing. No retrogradely labeled neurons were observed when AAV-DIO-RG was not injected. Scale bar, 200μm. (B) Validation of TVA-dependence of RV-mediated monosynaptic retrograde tracing. No retrogradely labeled neurons were observed when AAV-DIO-TVA-EGFP was not injected. Scale bar, 200μm. (C) A linear relationship was detected between the number of starter and input neurons. (D) Whole-brain distribution of normalized inputs (>1%) of PV+ (green) and SOM+ (red) neurons. Categorize them into auditory-related and polymodal-related brain regions based on brain functions. Data= means± S.E.M. (E) Normalized inputs of IC^SOM+^ and IC^PV+^ neurons from auditory-related and polymodal-related brain regions. N=4 mice per group, two-way ANOVA with Geisser-Greenhouse correction, ***p<0.001, ****p<0.0001. Data are means± S.E.M.

**supplement Figure 4.**
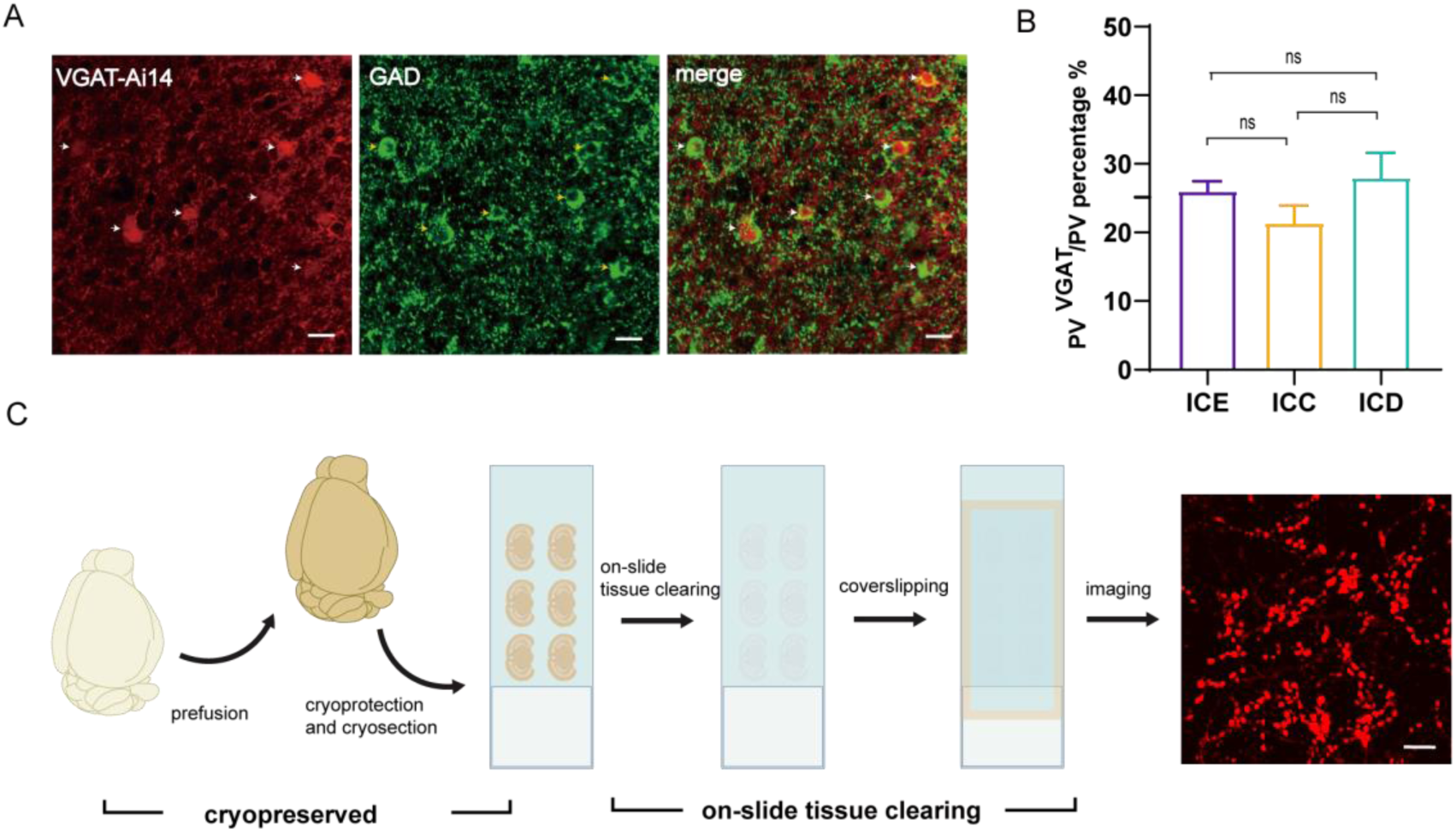
Expanded data to Figure 4. Anatomical characteristics of PV+ and SOM+ neurons in the IC. (A) Confocal 20× images showing VGAT neurons (red, white arrows, left), GAD65/67 staining (green, yellow arrows, middle), and an overlay (right). VGAT and GAD are co-expressed in the IC. Scale bars, 10μm. (B) Percentage of PV neurons co-labeled with VGAT (PV^VGAT^) among the total PV+ neurons in individual IC subdivision (ICE, ICC and ICD). n= 12 slices from N=3 mice; One-way ANOVA with post-hoc, p>0.5. (C) The procedure of rapid and non-scaling on-slide tissue clearing.

